# Spermidine enhances metabolic flexibility and attenuates inflammation associated with ageing in farmed Atlantic salmon

**DOI:** 10.64898/2026.03.13.711610

**Authors:** Kanchan Phadwal, Dominic Kurian, Jennifer Haggarty, Herve Migaud, Victoria Nicheva, James Dick, Muhammad Khalid F Salamat, Phillip D Whitfield, Chessor Matthew, Nicholas M. Wade, Mónica B Betancor, Daniel J. Macqueen

## Abstract

Metabolic ageing and associated changes in lipid mobilisation have been most heavily studied in humans and model taxa, yet remain poorly understood in farmed animals, with potentially important uncharacterised implications for health and welfare outcomes in food production systems. Here, we study both processes in domesticated Atlantic salmon (*Salmo salar*), the world’s most commercially valuable farmed fish, comparing three stages of aquaculture production. Our sampling captures a key life-cycle change where juvenile fish transition from freshwater into seawater (smoltification), followed by the ongoing ageing process during a final period of growth in seawater. Integrating lipidomics and proteomics of visceral adipose and skeletal muscle tissue, we firstly identified a metabolic-ageing profile akin to that observed in humans, which was distinct from lipid-associated remodelling associated with earlier smoltification. This was marked by impaired triglyceride storage, dysfunctional autophagy-lysosomal pathways, inflammation, fibrosis and reduced pathogen clearance pathways in visceral adipose tissue. In skeletal muscle, ageing was accompanied by reduced metabolic flexibility together with triglyceride and fatty acid accumulation, depletion of phospholipids, and a reduction in free fatty acids required for ATP production. We go on to provide experimental *in vivo* evidence that dietary spermidine supplementation suppresses adipose inflammation and reverses age-associated metabolic flexibility by re-establishing the buffering role of adipose tissue and enhancing fatty acid metabolism in skeletal muscle. Importantly, spermidine appears to reprogram lipid flux to counter metabolic ageing. As farmed Atlantic salmon exhibit key features of metabolic ageing observed in humans that appear linked to its recent domestication history, this species offers a novel model for ageing related studies of vertebrate metabolism.

## Introduction

Ageing is a degenerative physiological process that increases vulnerability to disease. Core hallmarks include genomic instability, telomere shortening, epigenetic alterations, loss of proteostasis, metabolic dysregulation, and chronic inflammation (López-Otín et al. 2013). While these processes are highly characterized in mammals (including humans) and model organisms, they remain largely unexplored in non-model species, notably including species farmed for human consumption. This represents a significant knowledge gap, firstly in terms of understanding the diversity of ageing mechanisms present in nature, but also how these processes impact the health, robustness and welfare of farmed animals, which may have knock-on effects for the health of human consumers.

Atlantic salmon (*Salmo salar,* hereafter: salmon) is one of the most important farmed fish species globally (Houston and Macqueen 2019). In common with other intensively farmed animals, salmon have been strongly domesticated - for instance through artificial rearing conditions, selective breeding, and high-lipid diets. Consequently, farmed salmon show highly accelerated growth compared to wild-type ancestors, shortening the production time required for them to reach advanced stages of the life-cycle (Jonsson, Jonsson, and Finstad 2013; McLennan et al. 2016; Åsheim et al. 2023).

High-fat diets have emerged as a potentially key driver of accelerated metabolic ageing via mitochondrial and lysosomal dysfunction, leading to chronic inflammation in humans and mice (Heijden et al. 2015; van der Heijden et al. 2015; Carmona-Gutierrez et al. 2016; Balasubramanian et al. 2024; Zhang et al. 2025). Modern farmed salmon diets are rich in vegetable oils, which are high in saturated fatty acids (SFA) and monounsaturated fatty acids (MUFA), as well as ω-6 polyunsaturated fats (PUFA) (Olsen and Skjervold 1995; Katan et al. 2021; Molversmyr et al. 2022). These oils, primarily from soybean (rich in linoleic acid, 18:2ω-6) and rapeseed (high in oleic acid, 18:1ω-9 and linoleic acid), have been used to address sustainability issues surrounding finite marine fish oils rich in ω-3 PUFA such as eicosapentaenoic acid (EPA, 20:5ω -3) and docosahexaenoic acid (DHA, 22:6ω-3) (Sprague, Dick, and Tocher 2016). However, this dietary shift has been associated with excessive lipid accumulation in salmon visceral tissues, with health impacts on the fish, as well as reducing the health-promoting ω-3 EPA and DHA content of fillets consumed by humans (Bell et al. 1998; Weihe et al. 2019; Molversmyr et al. 2022).

Lipid accumulation in tissues is a hallmark of metabolic ageing in vertebrates (Lee, Olson, and Evans 2003; Chung 2021). Normally, surplus lipids are stored as triglycerides (TGs) in white adipose tissue. However, when adipose tissue becomes dysfunctional or overloaded, excess lipids accumulate ectopically in the skeletal muscle, liver, and kidney (Chung 2021). Moreover, chronic dysfunction in white adipose tissue promotes lipotoxicity in peripheral tissues, driving systemic inflammation, metabolic disruption, and accelerated ageing (Ou et al. 2022). In skeletal muscle, excess lipid deposition is closely linked to reduced metabolic flexibility – leading to a diminished ability to switch efficiently between fuels. This is thought to arise not only from increased intramyocellular TG storage, but also from the accumulation of lipid intermediates that impair mitochondrial substrate handling, promoting incomplete β-oxidation, energetic inefficiency, and progressive functional decline with age (Ukropcova et al. 2005; Bennett and Sato 2023).

The salmon farming industry has recently seen a rise in mortality (Norwegian Veterinary Reports. 2026), concentrated in the final seawater phase of production where fish are grown until harvested for human consumption, reaching sizes typical of advanced life-stages in nature. We hypothesize that such late-stage mortality is driven in part by metabolic ageing, expedited by production intensification from domestication and high fat diets, leading to systemic inflammation, which has been shown to compromise immune resilience and increase susceptibility to disease (Ou et al. 2022; Dai et al. 2023). To test this hypothesis, we collected visceral adipose tissue (mesenteric, hereafter: ‘VAT’), a fat depot and secondary lymphoid organ in teleosts (Simón et al. 2022) and fast-twitch skeletal muscle (hereafter: ‘SM’), also a fat depot, from salmon at three life-stages ranging from juvenile to advanced adult seawater stages (**Figure 1**). In addition to providing insights into metabolic ageing during the seawater growth phase, our sampling captures a juvenile life-history transition called smoltification, which is accompanied by radical changes in metabolism and energy usage, including for lipids (Sheridan, 1989; Maxime 2002; Handeland et al. 2003; Björnsson, Stefansson, and McCormick 2011). High-resolution lipidomics and proteomics were used to reveal a global signature of metabolic ageing akin to that reported in human and rodent models, while also capturing fatty-acid and lipid-metabolism shifts during smoltification. To experimentally test the reversibility of metabolic ageing, we supplemented salmon diets with the polyamine spermidine, a compound with established anti-ageing effects in mammals (Madeo, Carmona-Gutierrez, et al. 2018). Our results provide insights into the conserved nature of metabolic ageing in vertebrates, offer a novel dietary strategy to reduce associated potentially pathological impacts on fish health, and highlight the value of salmon as a model for ageing research.

**Figure 1.**
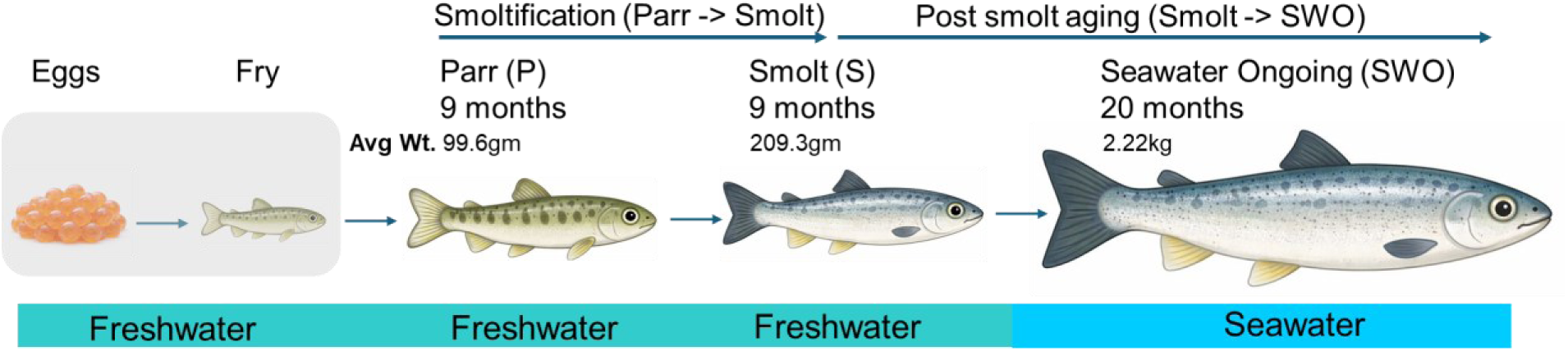
Summary of salmon life stage transitions sampled for lipidomic and proteomic analysis of skeletal muscle and visceral adipose tissue. The boxed grey early life stages were not sampled, and were included to illustrate all major stages of aquaculture production.

## Results

### The muscle and adipose lipidome is reshaped across salmon life-stages

Different phases of the salmon life-cycle have been associated with substantial metabolic and immunological remodelling, which are thought to represent adaptations to distinct environments (Sheridan 1989; Handeland et al. 2003; Björnsson et al. 2011; Johansson et al. 2016; van Muilekom et al. 2024). To investigate if different salmon life-stages are accompanied by changes in the lipid storage pool, we compared lipidomic profiles for SM and VAT between parr, smolt, and seawater ongoing (SWO) stage farmed salmon (**Figure 1**).

Across tissues, triglycerides (TGs) were the dominant lipid class, with SM enriched for phospholipids (phosphatidylethanolamine, PE; phosphatidylcholine, PC), whereas VAT predominantly accumulated TGs (**Table 1; Supplementary Figure 1**). Principal component analysis (PCA) of the lipidome separated fish by stage in both tissues (**Figure 2a-b**). For interpretation of neutral lipid changes, TGs are grouped by total carbon number (CN): medium-chain TGs (mcTGs; CN 40-48), long-chain TGs (lcTGs; CN 50-58) and very-long-chain TGs (vlcTGs; CN > 60). Lipid species that differed across stages are summarised in **Table 1**.

**Table 1.**
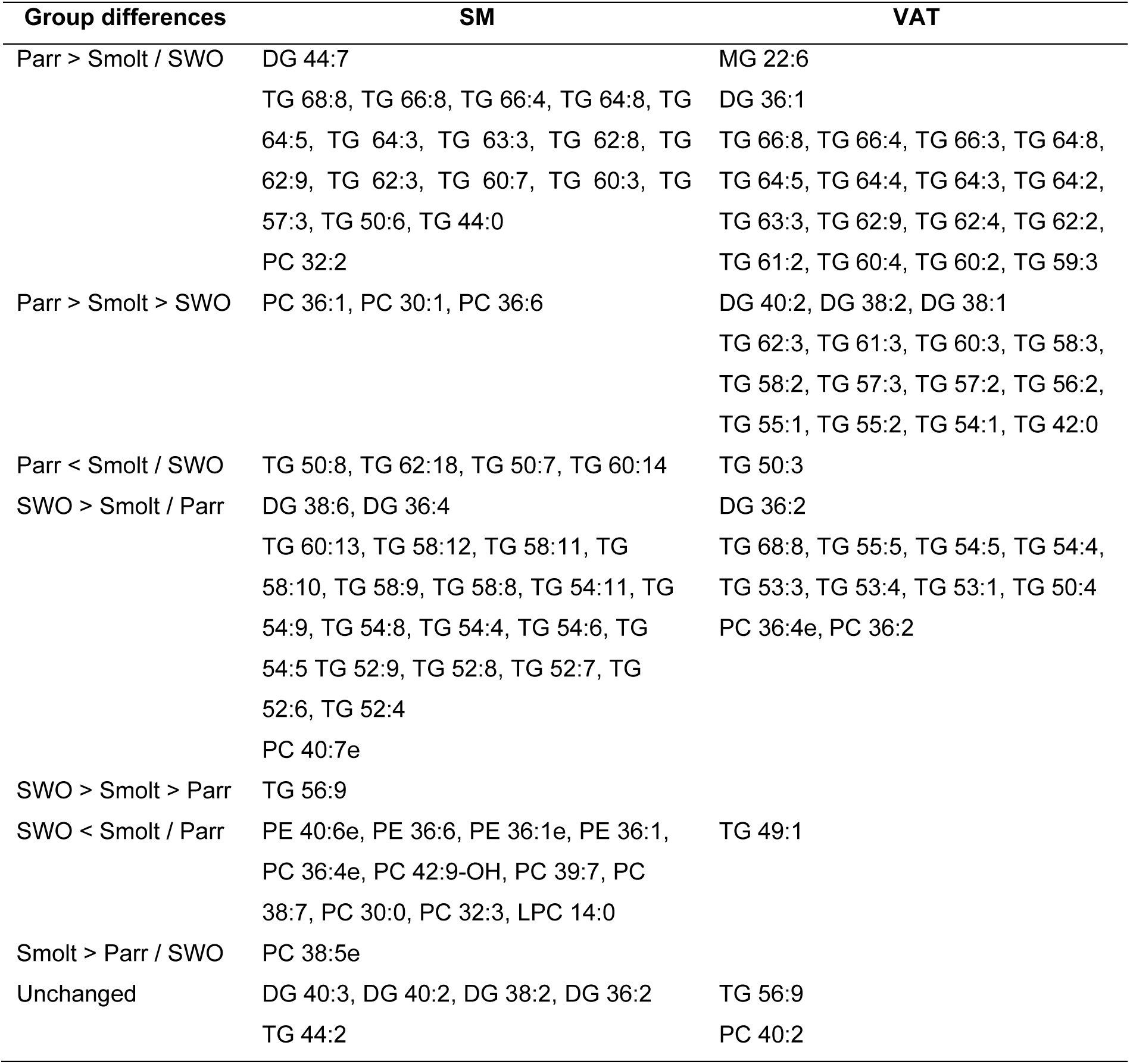
Changes in lipidome across three life-stages in SM and VAT.

| Group differences | SM | VAT |
| --- | --- | --- |
| Parr > Smolt / SWO | DG 44:7<br>TG 68:8, TG 66:8, TG 66:4, TG 64:8, TG 64:5, TG 64:3, TG 63:3, TG 62:8, TG 62:9, TG 62:3, TG 60:7, TG 60:3, TG 57:3, TG 50:6, TG 44:0<br>PC 32:2 | MG 22:6<br>DG 36:1<br>TG 66:8, TG 66:4, TG 66:3, TG 64:8, TG 64:5, TG 64:4, TG 64:3, TG 64:2, TG 63:3, TG 62:9, TG 62:4, TG 62:2, TG 61:2, TG 60:4, TG 60:2, TG 59:3 |
| Parr > Smolt > SWO | PC 36:1, PC 30:1, PC 36:6 | DG 40:2, DG 38:2, DG 38:1<br>TG 62:3, TG 61:3, TG 60:3, TG 58:3, TG 58:2, TG 57:3, TG 57:2, TG 56:2, TG 55:1, TG 55:2, TG 54:1, TG 42:0 |
| Parr < Smolt / SWO | TG 50:8, TG 62:18, TG 50:7, TG 60:14 | TG 50:3 |
| SWO > Smolt / Parr | DG 38:6, DG 36:4<br>TG 60:13, TG 58:12, TG 58:11, TG 58:10, TG 58:9, TG 58:8, TG 54:11, TG 54:9, TG 54:8, TG 54:4, TG 54:6, TG 54:5 TG 52:9, TG 52:8, TG 52:7, TG 52:6, TG 52:4<br>PC 40:7e | DG 36:2<br>TG 68:8, TG 55:5, TG 54:5, TG 54:4, TG 53:3, TG 53:4, TG 53:1, TG 50:4<br>PC 36:4e, PC 36:2 |
| SWO > Smolt > Parr | TG 56:9 |  |
| SWO < Smolt / Parr | PE 40:6e, PE 36:6, PE 36:1e, PE 36:1, PC 36:4e, PC 42:9-OH, PC 39:7, PC 38:7, PC 30:0, PC 32:3, LPC 14:0 | TG 49:1 |
| Smolt > Parr / SWO | PC 38:5e |  |
| Unchanged | DG 40:3, DG 40:2, DG 38:2, DG 36:2<br>TG 44:2 | TG 56:9<br>PC 40:2 |

**Figure 2.**
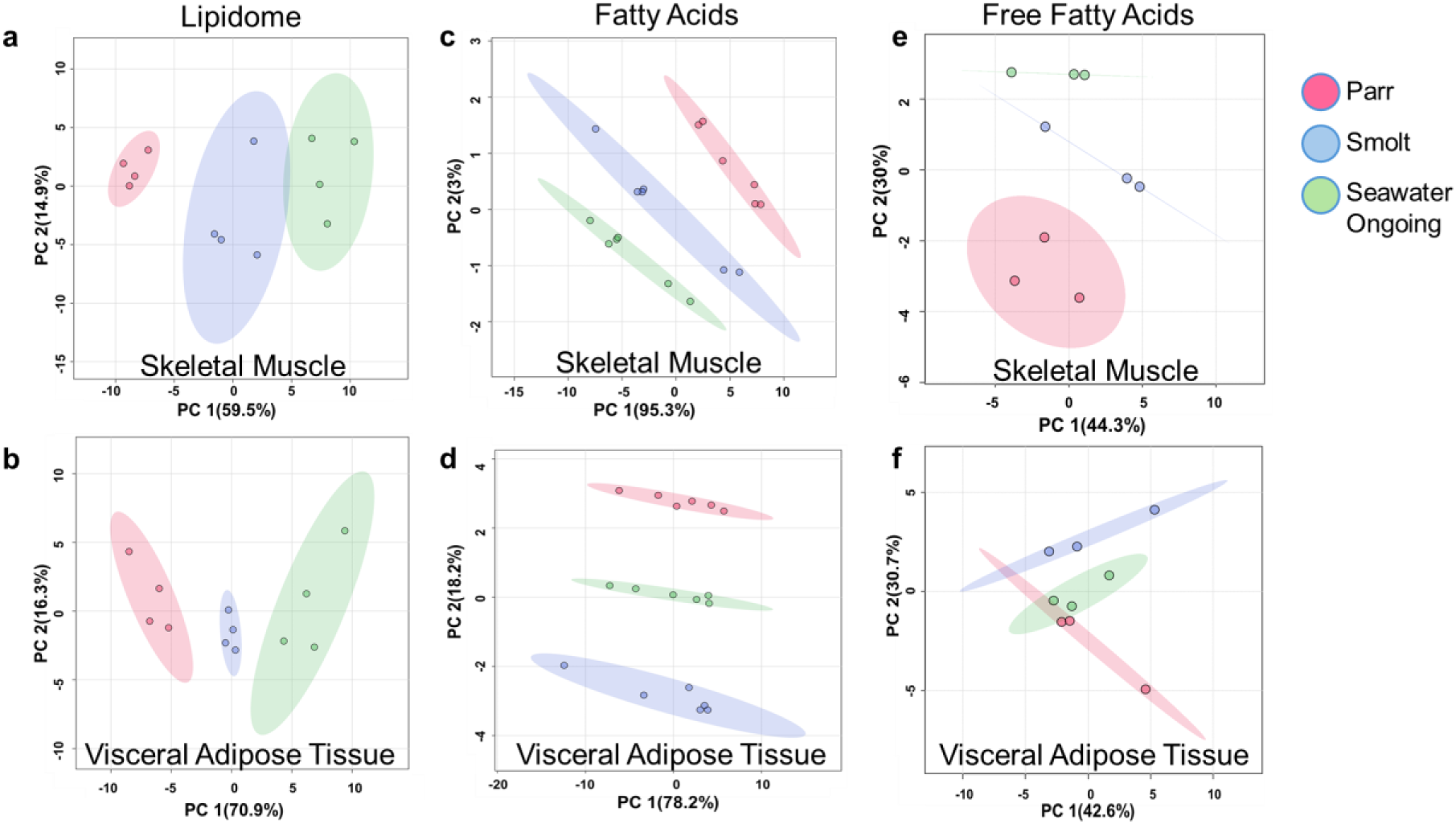
Principal component analysis of lipid profiling datasets across the sampled life stages. Lipidomics for SM (n=4) (**a**) and VAT (n=4) (**b**). Fatty acid analysis for SM (n=6) (**c**) and VAT (n=6) (**d**). Free fatty acid analysis for SM (n=3) (e) and VAT (n=3) (f).

In SM, smoltification (from parr to smolt stage) and post-smolt ageing (from smolt to SWO stage) were characterised by reduced abundance of vlcTGs and increased abundance of lcTGs (**Table 1, Supplementary Figure 1a**). Changes during smoltification and post-smolt ageing were distributed across TG species (**Table 1**). Multiple phospholipid types declined with progression to SWO, including PCs and PEs. PC 38:5e increased during smoltification, but decreased during post-smolt ageing, while PC 36:1, PC 30:1 and PC 36:6 showed a gradual decline across life stages. Several other PCs (PC 36:4e, PC 42:9-OH, PC 39:7, PC 38:7, PC 30:0 and PC 32:3), PEs (PE 40:6e, PE 36:6, PE 36:1e and PE 36:1) and LPC 14:0 were lower in SWO than parr and/or smolt (**Table 1**, **Supplementary Figure 1b**).

In VAT, parr showed higher abundance of several vlcTGs compared with smolt and SWO (**Table 1, Supplementary Figure 1c**). Multiple DGs and TGs (including mcTGs, lcTGs and vlcTGs) decreased progressively across stages (parr > smolt > SWO). In contrast, a subset of mcTGs and lcTGs increased in SWO compared with parr and/or smolt (SWO > smolt / parr). Among phospholipids, PC 36:4e and PC 36:2 were higher in SWO (**Table 1**).

### Ageing salmon muscle shows greater accumulation of fatty acids than adipose

Using a GC/MS-based approach, we quantified total fatty acids (FAs) - the building blocks of complex lipids - in SM and VAT from parr, smolt and SWO stage salmon (n=6 per group) (**Table 2**). PCA for total FA profiles separated fish by stage in both tissues (**Figure 2c-d)**.

**Table 2.** FA profile changes across the three life-stages in SM and VAT.

| **Group differences** | **SM** | **VAT** |
| --- | --- | --- |
| Parr > Smolt > SWO |  | **MUFA:** FA22:1n-11, FA20:1n-11 |
| Parr < Smolt / SWO | **MUFA:** FA22:1n-9, FA20:1n-7,<br>FA18:1n-7, FA16:1n-7.<br>**PUFA- Omega 6:** FA22:5n-6,<br>FA20:4n-6, FA20:2n-6, FA18:3n-6,<br>FA18:2n-6,<br>**PUFA- Omega 3:** FA22:5n-3,<br>FA21:5n-3, FA20:5n-3, FA20:4n-3,<br>FA18:4n-3, FA18:3n-3<br>**SFA:** FA22:0, FA20:0, FA18:0, FA<br>16:0, FA15:0, FA14:0 |  |
| SWO > Smolt / Parr | **MUFA:** FA18:1n-9, FA16:1n-9<br>**PUFA- Omega 6:** FA20:3n-6<br>**SFA:** FA24:0 |  |
| SWO > Smolt > Parr | **MUFA:** FA20:1n-9<br>**PUFA- Omega 3:** FA18:3n-3,<br>FA20:3n-3<br>**PUFA- Omega 6:** FA18:2n-6,<br>FA20:2n-6, FA22:5n-6 |  |
| Unchanged | **MUFA:** FA22:1n-11, FA20:1n-11,<br>FA24:1n-9<br>**PUFA- Omega 3:** FA22:6n-3 | **MUFA:** FA22:1n-9, FA20:1n-9,<br>FA 18:1n-9, FA16:1n-9<br>FA20:1n-7, FA18:1n-7, FA<br>16:1n-7<br>**PUFA- Omega 6:** FA22:5n-6,<br>FA 20:2n-6, FA 20:3n-6, FA<br>18:3n-6, FA18:2n-6, FA 20:4n-6,<br>**PUFA- Omega 3:** FA21:5n-3,<br>FA 20:5n-3, FA22:6n-3,<br>FA22:5n-3, FA20:4n-3, FA<br>20:3n-3, FA18:4n-3, FA18:3n-3<br>**SFA:** FA24:0, FA22:0, FA20:0,<br>FA18:0, FA16:0, FA15:0, FA<br>14:0 |

In SM, most FAs were higher in smolt and SWO than parr (**Table 2, Supplementary Figure 2a**), spanning MUFAs, omega-3 and omega-6 PUFAs, and saturated FAs. A subset of FAs were higher in SWO than parr and smolt, including MUFAs 20:1n-9, 18:1n-9 and FA16:1n-9, the n-6 PUFA 20:3n-6, and saturated 24:0 (**Table 2**). MUFA: 20:1n-9, n-3 PUFA: 18:3n-3, 20:3n-3 and n-6 PUFA: 18:2n-6, 20:2n-6, 22:5n-6 FAs increased progressively across stages (SWO > smolt > parr). Several FAs remained unchanged across stages, including the MUFAs 22:1n-11, 20:1n-11 and 24:1n-9, and docosahexaenoic acid (n-3 22:6n-3) (**Table 2**). In VAT, total FA levels remained similar, with FA 22:1n-11 and FA 20:1n-11 showing a decline at each successive life stage (parr > smolt > SWO) (**Table 2, Supplementary Figure 2b**).

Next, we examined if changes in total FAs were reflected in the free fatty acid (FFA) pool. PCA of FFAs separated fish by stage in both tissues (**Figure 2e-f**). In SM, FFAs provide a substrate for mitochondrial beta-oxidation and ATP generation to support growth and swimming activity (Stubhaug, Frøyland, and Torstensen 2005), whereas VAT serves as a major TG storage depot that mobilises FFAs during increased energy demand (Bou et al. 2020). In SM, parr showed higher abundance of two MUFA FFAs (22:1n-9 and 20:1n-9) compared with smolt and SWO, whereas the omega-3 PUFA FA18:3n-3 was higher in SWO than parr and smolt (**Table 3, Supplementary Figure 3a**). Beyond these differences, most FFAs were unchanged across stages. In VAT, no significant stage-associated differences were detected in the FFA pool (**Table 3, Supplementary Figure 3b**).

**Table 3.**
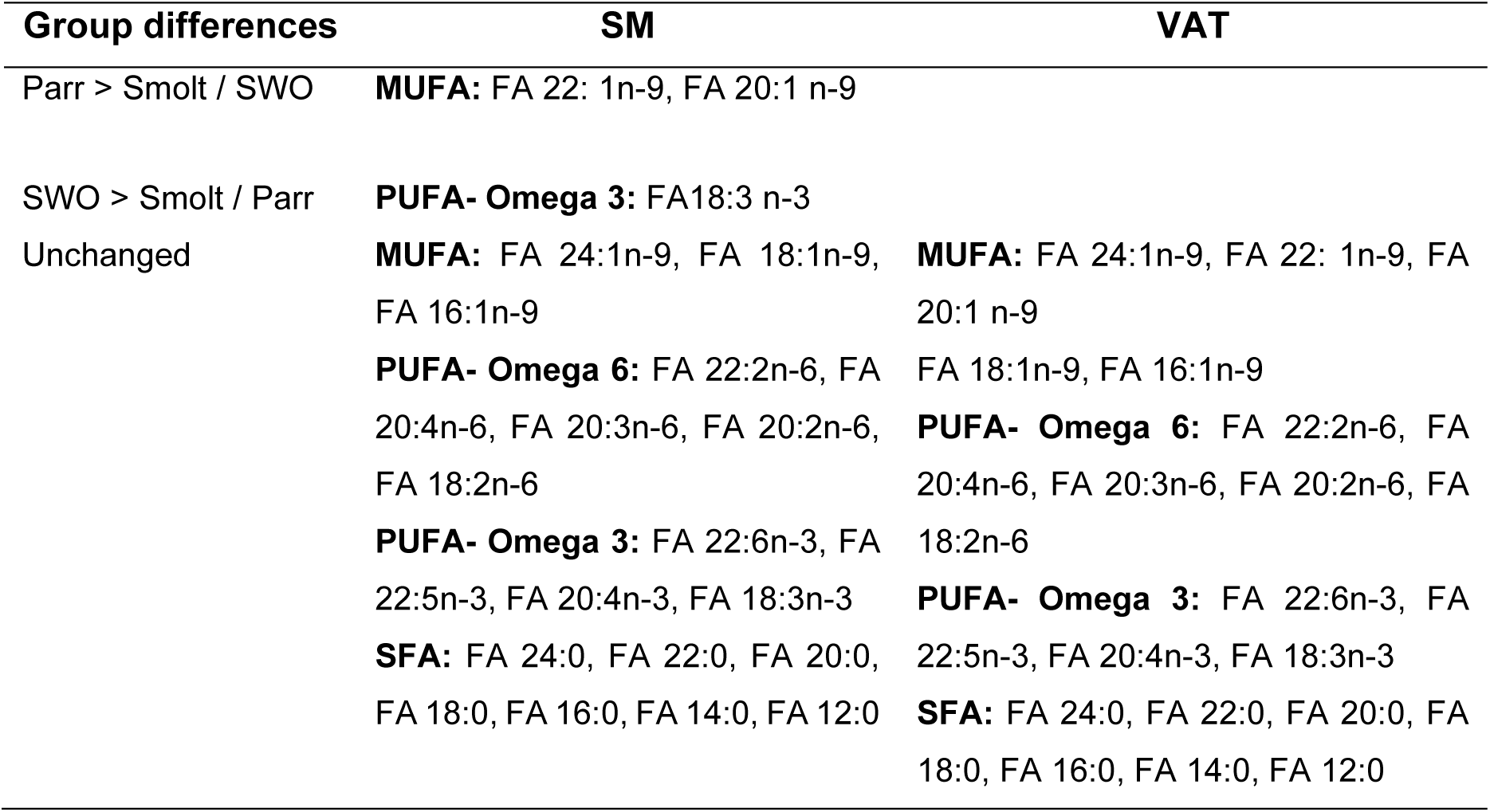
Changes in FFA profile across the three life stages in SM and VAT.

### Lipid overload metabolic phenotype in muscle during seawater ageing

Despite lipid accumulation in SM, FFAs did not rise in the SWO stage, motivating further interrogation of oxidative capacity during smoltification and ageing through proteomic profiling of SM and VAT in parr, smolt, and SWO (n=5 per group).

2,623 proteins were identified in SM, with 197 showing differences in abundance (*p* < 0.05) between parr and smolt and 161 between smolt and SWO (**Supplementary Data 1a-b**; **Figure 3a-b**). GO terms enriched in SM related to carbohydrate metabolism, including carbon metabolism, glycolysis/gluconeogenesis, the pentose phosphate pathway, and starch and sucrose metabolism (**Supplementary Figure 4a**). However, this enrichment was specific to smoltification (parr vs. smolt). Compared to parr, the smolt stage showed upregulation of carbohydrate-metabolism proteins, including fructose-1,6-bisphosphatase 2 (FBP2), aldolase A (ALDOA), pyruvate kinase M, isoform B (PKMB), and β-enolase (ENO3), indicating a shift toward carbohydrate use (**Figure 3a**). Acyl-CoA synthetase long-chain family member 1a and 1B (ACSL1A/B) were upregulated in parr compared to both smolt and SWO SM, indicating efficient fatty-acid based ATP production from lcTGs and vlcTGs (**Figure 3a**). Parr further showed upregulation of markers of glycolysis (PFKM, GPI1) and oxidative phosphorylation (COX6A2, COX6B2 – cytochrome c oxidase subunits (Complex IV; aerobic capacity) and NDUFA5 – Complex I subunit (electron transport chain) (**Supplementary Figure 5**) compared to smolt and SWO; again, suggestive of active ATP generation pathways.

**Figure 3.**
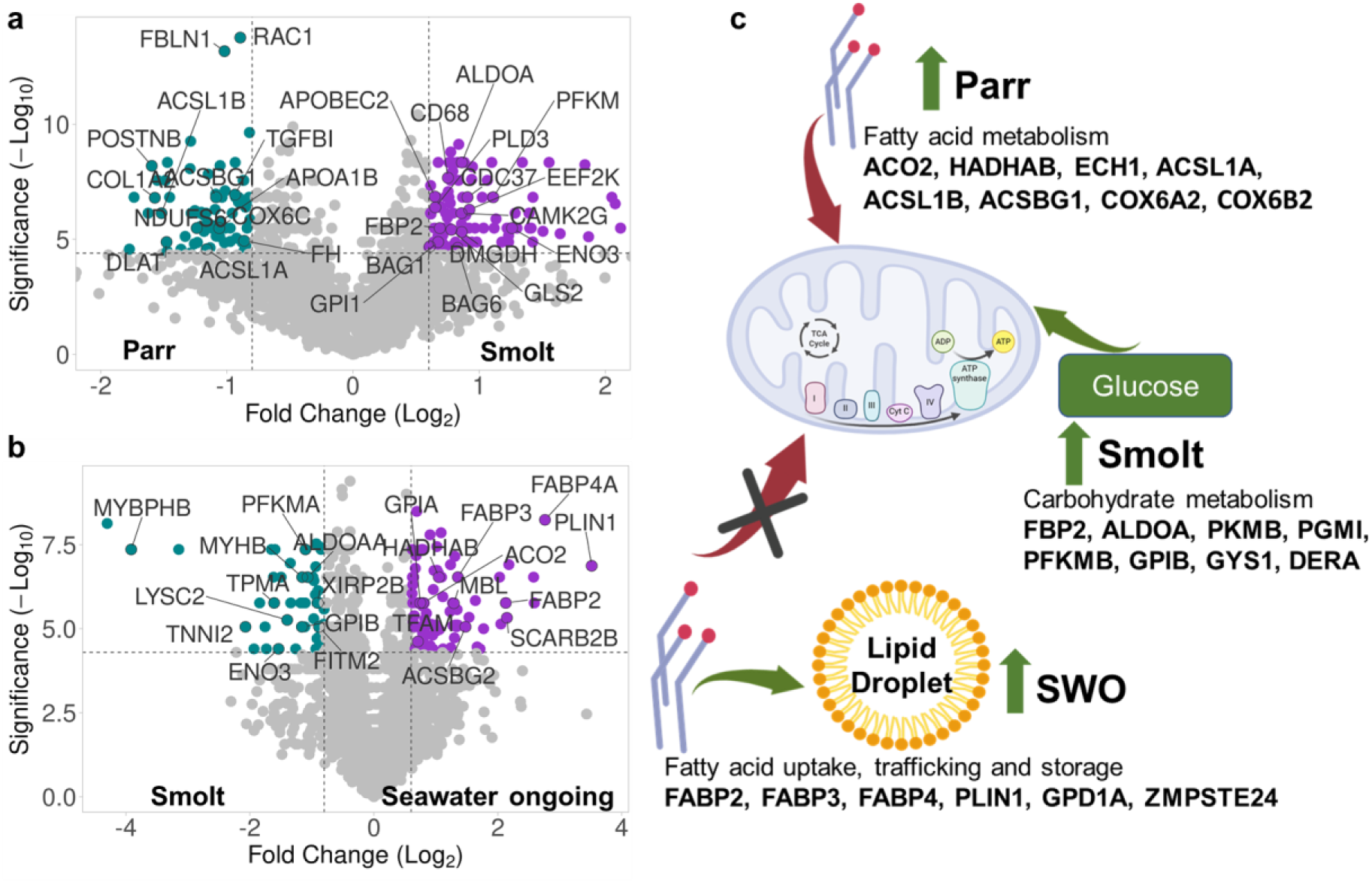
Proteomics reveals a lipid overload-driven metabolic ageing phenotype in the skeletal muscle of seawater ongoing salmon. Results shown for: parr (n=6), smolt (n=5), and SWO (n=5). Volcano plots show up (purple dots) and downregulated (teal dots) proteins between (**a**) parr and smolt. (**b**) smolt and seawater ongoing (SWO). (**c**) Summary of metabolic remodelling in skeletal muscle as ontogeny proceeds (parr → smolt → SWO). Arrows indicate inferred substrate flux through mitochondrial metabolism, and up-arrows denote stage-associated increases in the abundance of key proteins.

Compared to parr and smolt, SWO salmon did not show upregulation of any carbohydrate or fatty acid catabolic pathways. Instead, post-smolt ageing (smolt -> SWO) was associated with increased abundance of proteins involved in lipid uptake, trafficking, and storage; such as fatty acid binding protein 2, 3 and 4 (FAB2/3/4), glycerol-3-phosphate dehydrogenase 1A (GPD1A), and perilipin 1 (PLIN1), suggesting enhanced lipid influx and sequestration rather than oxidation (**Figure 3b**).

Key markers supporting fatty acid breakdown (parr), glycolysis (smolt) and fatty acid storage (SWO) are summarised in **Figure 3c**. Abundance changes in proteins contributing to metabolic signatures across life-stages are presented in **Supplementary Figure 5a-q**.

### Declining autophagy-lysosomal pathways and signatures of fibrosis and inflammation in VAT during seawater ageing

In VAT, we identified 5,342 proteins (**Supplementary Data 2a**). Of these, 48 were differentially abundant (p < 0.05) between parr and smolt (**Figure 4a**) and 193 between smolt and SWO (**Figure 4b; Supplementary Data 2b**).

**Figure 4.**
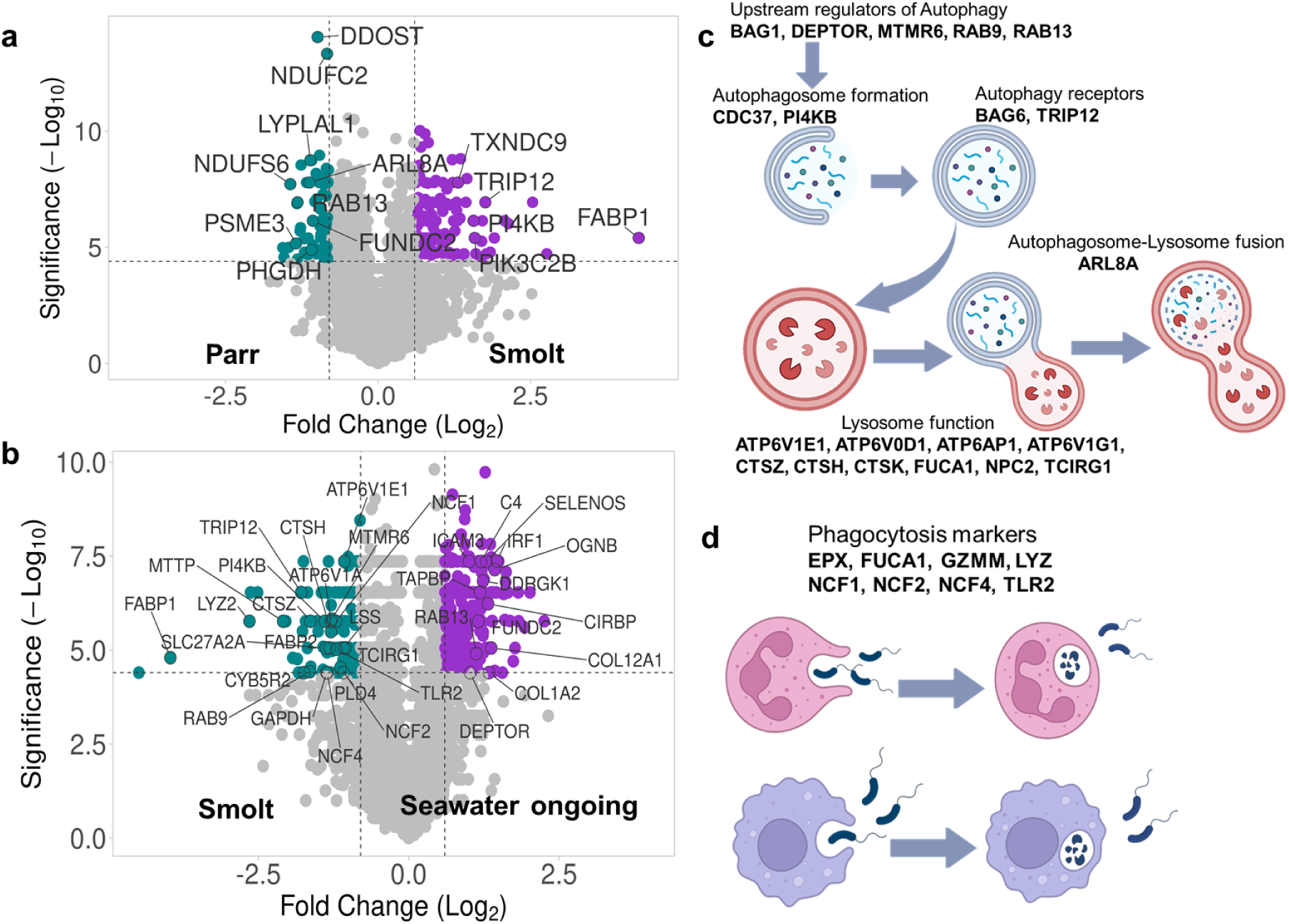
Proteomics reveals a decline in the autophagy-lysosomal pathway with a signature of fibrosis and inflammation in visceral adipose tissue of seawater ongoing salmon. Results are shown for: parr (n=5), smolt (n=5), and seawater ongoing (n=5). Volcano plots show both up (purple dots) and downregulated (teal dots) proteins between (**a**) parr and smolt, (**b**) smolt and seawater ongoing. Also shown is a summary of autophagy-lysosome (**c**) and phagosome (**d**) pathways downregulated in seawater ongoing, including key proteins.

The main GO terms enriched in VAT were related to lysosomes, phagosomes, metabolic pathways (including eicosanoid / arachidonic-acid oxylipin metabolism and carbohydrate metabolism), ECM - receptor interaction and cytoskeleton (**Supplementary Figure 4b-c**). However, we did not find any GO term enrichment during smoltification (parr -> smolt).

Compared to parr and smolt, the SWO proteome revealed a marked decline in proteins required for lysosomal function, including multiple lysosomal hydrolases (cathepsin Z; CTSZ, cathepsin H; CTSH, niemann–pick disease, type c2 protein; NPC2, phospholipase D4; PLD4, alpha-l-fucosidase 1; FUCA1, T cell immune regulator 1; TCIRG1), lysosomal acidification pumps (ATPase H⁺ transporting v1 subunit e1; ATP6V1E1, ATPase H⁺ transporting V0 subunit d1: ATP6V0D1), and proteins required for lysosome–autophagosome fusion (ADP-ribosylation factor–like protein 8A; ARL8A). Further, several autophagy regulators (myotubularin-related protein 6; MTMR6, ras-related protein Rab-9: RAB9), proteins necessary for autophagosome formation (phosphatidylinositol 4-kinase beta; PI4KB), and regulation of autophagy cargo–recognition factors (thyroid hormone receptor interactor 12; TRIP12) were specifically downregulated in SWO (**Figure 4c, Supplementary Figure 6a-g, Supplementary Figure 7a-f**).

Compared to parr and smolt, SWO also showed decreased abundance of proteins central to phagocytosis and innate immune defence, including eosinophil peroxidase (EPX), alpha-l-fucosidase 1 (FUCA1), granzyme m (GZMM), lysozyme 2 (LYZ2), neutrophil cytosolic factor 1, 2 and 4 (NCF1/2/4) and the pathogen recognition receptor toll-like receptor 2 (TLR2) (**Figure 4d & Supplementary Figure 6h-o**).

On the other hand, SWO displayed a strong increase in abundance of proteins associated with adipose inflammation clustered under enriched ECM receptor interaction and cytoskeleton pathways: complement component 4; C4, serpin family g member 1; SERPING1, intercellular adhesion molecule 3; ICAM3, tapasin; TAPBP, interferon regulatory factor 1 and 3; IRF1/3, osteoglycin; OGNB, proline/arginine-rich end leucine-rich repeat protein; PRELP, along with fibrosis markers cytoskeletal markers collagen type xii alpha 1 chain (COL12A1), collagen type i alpha 2 chain (COL1A2), decorin (DCN) and laminin subunit alpha 4 (LAMA4) (**Figure 4a-b, Supplementary Figure 7g-o)**. Comparative plots of these key pathway components across life stages are provided in **Supplementary Figures 6-7.**

In comparison to smolt, SWO showed decreased abundance of proteins involved in eicosanoid / arachidonic-acid oxylipin metabolism within the metabolic pathway cluster, including prostaglandin-endoperoxide synthase 1 (PTGS1) (commonly called cyclooxygenase-1, COX-1), arachidonate 8-lipoxygenase (ALOX8), ATP-binding cassette subfamily c member 4 (ABCC4) and carbohydrate metabolism glucokinase (GCK) (also called hexokinase IV), glyceraldehyde-3-phosphate dehydrogenase (GAPDH) and pyruvate kinase (PKLR) (**Supplementary Figure 4c and Supplementary Figure 6p-u**).

### Spermidine activates autophagy and attenuates metabolic ageing phenotypes

We next asked if the metabolic ageing phenotype inferred in SWO can be reversed using the natural polyamine spermidine. Oral spermidine supplementation extends lifespan in multiple species (Eisenberg et al. 2009), while mitigating high-fat diet–induced obesity and metabolic dysfunction, and reducing inflammation in VAT (Liao et al. 2021; Wang et al. 2021). These effects are primarily attributed to autophagy induction ( Madeo, Bauer, et al. 2018; Green 2019).

We compared SWO salmon fed a control (soy oil-enriched) diet with a group fed the same diet supplemented with 5 mg/kg spermidine (after: Zhang et al. 2017, 2021; Guo et al. 2022) (**Supplementary Figure 8**). There were no differences in growth parameters between groups after 5 weeks (**Supplementary Figure 8e-i**).

Using lipidomics, we identified four vlcTGs in SM reduced by spermidine (TG 70:8, TG 66:3, TG 65:3 and TG 63:3), in parallel to an increase in several phospholipids (including PC 38:6, LPC 18:2 and LPC 16:0) (**Figure 5a**). VAT showed a broader range of lipid subclasses, with the spermidine group showing increased abundance of diverse lcTGs (TG 48:3, TG 52:4, TG 53:4, TG 54:0, TG 54:3, TG 55:4, TG 56:3) and phospholipids (PS, PI, PE, PC and LPC) (**Figure 5b**).

**Figure 5.**
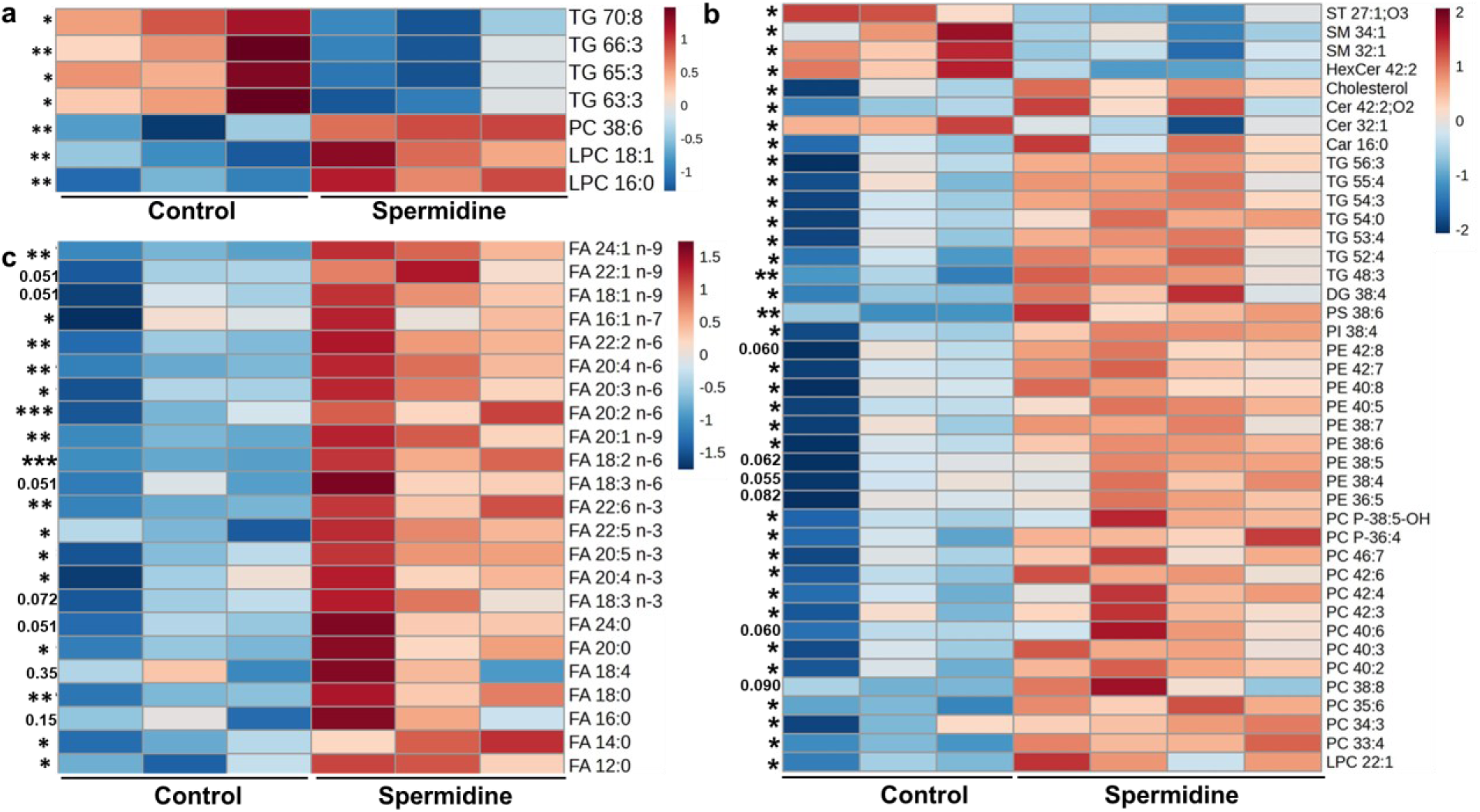
Spermidine supplementation reverses age-related defects in lipid storage. Heatmaps are shown to illustrate changes in (**a**) the skeletal muscle lipidome (n=3), (**b**) the visceral adipose tissue lipidome (n = 3/4), and (**c**) the skeletal muscle free fatty acids (n = 3). Data shown are mean +/− SD (*\* p < 0.05; ** p < 0.01; *** p < 0.001*). Non-significant values are reported as exact p-values.

The spermidine group showed reduced levels of lipotoxic lipids in VAT, including ceramides Cer(d18:1/14:0), sphingomyelins SM(d18:1/14:0), and oxysterols ST27:1; O3, which have been linked to obesity, insulin resistance, and inflammation in mammals (Brusatori et al. 2022; Guillemot-Legris et al. 2016; Haley et al. 2017; Montgomery et al. 2017) (**Figure 5b**). In addition, we identified an increase in palmitoyl-L-carnitine (Car 16:0), a metabolite that promotes β-oxidation by transporting long-chain fatty acids such as palmitic acid across mitochondrial membranes (Uner et al. 2023) (**Figure 5b**). Cholesterol, which is part of adipocyte membranes and required for adipocyte function also increased in VAT from the spermidine group (**Figure 5b**).

We did not identify any difference between FA abundance in SM or VAT comparing the spermidine and control diet groups. However, SM from the spermidine group showed an increase in several FFAs, including MUFAs and SFAs (**Figure 5c**). In contrast, no changes in FFA levels were observed in VAT, barring a decline in FA 22:1n-9 (data not shown).

As spermidine is known to enhance autophagic capacity, we next assessed its effects on various autophagy markers in SM. We measured the abundance of sequestosome-1 (p62/SQSTM1), a selective autophagy adaptor and marker for autophagic degradation (Bjørkøy et al. 2009), and autophagy-related protein 7 (ATG7), a core autophagy protein required for autophagosome formation (Cawthon et al. 2018). Abundance changes of p62 and Atg7 in SM indicate increased autophagic capacity in the spermidine group (**Figure 6a-d**). We also found that SM from the spermidine group showed an increase in carnitine palmitoyltransferase 1A (CPT1A) (**Figure 6e-f**), an enzyme required to catalyse the rate-limiting step in transporting long-chain fatty acids into mitochondria for β-oxidation (Bruce et al. 2009).

**Figure 6.**
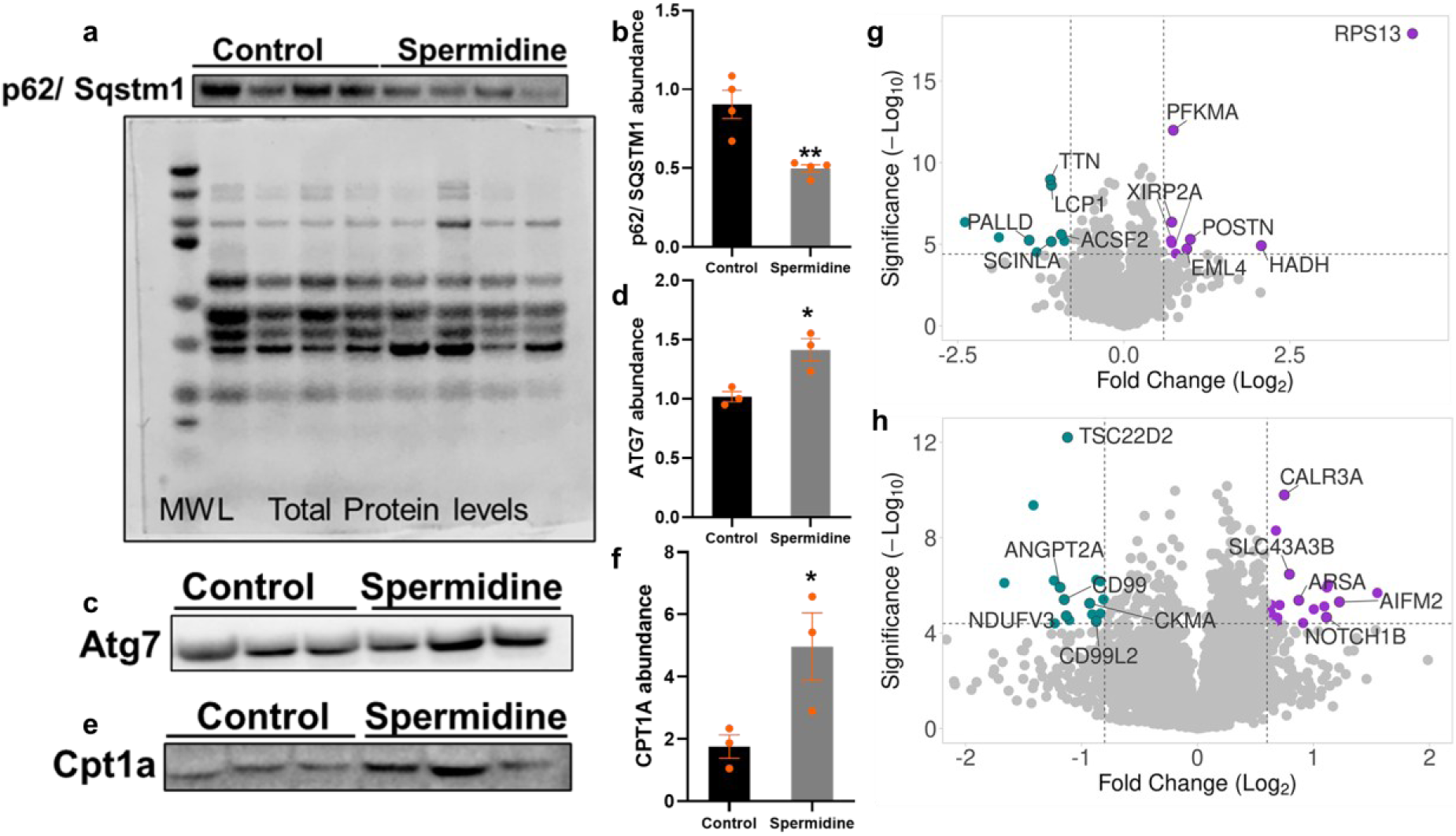
Spermidine supplementation enhances autophagy and promotes a metabolic shift in skeletal muscle while suppressing inflammation in visceral adipose tissue. Protein abundance changes inferred by immunoblot quantification for: (**a**-**b**) p62/ Sqstm1 (n=4) (**a**) also shows total protein levels used for normalisation), (**c-d**) Atg7 (n=3), and (**e-f**) Cpt1a (n=3). Data shown are mean +/− SD (* *p* < 0.05; ** *p* < 0.01). Proteomics data are shown as volcano plots highlighting proteins that are up-(purple dots) or down-regulated (teal dots) following spermidine supplementation in (**g**) skeletal muscle (n=3 per group) and (**h**) visceral adipose tissue (n=3 per group).

Using proteomics in SM, 2,454 proteins were detected, with 8 upregulated and 9 downregulated in the spermidine group (**Figure 6g; Supplementary Data 3a-b**). Proteins upregulated included those involved in translation (RPS13), energy metabolism—covering both glycolysis (phosphofructokinase muscle a-isoform; PFKMA) and mitochondrial β-oxidation (hydroxyacyl-CoA dehydrogenase; HADH), three paralogs of xin actin–binding repeat–containing protein 2a (XIRP2A) and periostin (POSTN). Conversely, spermidine downregulated lymphocyte cytosolic protein 1 (LCP1), titin (TTN), acyl-coA synthetase family member 2 (ACSF2) and palladin (PALLD) (**Figure 6g**). In VAT, 9,704 proteins were identified with 23 upregulated and 20 downregulated in the spermidine group (**Figure 6h**; **Supplementary Data 4a-b**). Proteins upregulated included solute carrier family 43 member 3 (Slc43a3), the lysosomal enzyme arylsulfatase A (ARSA), and apoptosis-inducing factor mitochondria associated 2 (AIFM2), while downregulating the inflammatory proteins Angiopoietin 2a (ANGPT2A), CD99 and CD99L2.

## Discussion

Our results demonstrate that ageing in farmed salmon is accompanied by lipid dysregulation and loss of metabolic flexibility in skeletal muscle (SM), alongside dysfunctional lysosomal and phagocytotic functions in parallel to inflammation and fibrosis in visceral adipose tissue (VAT). These changes map closely to metabolic ageing phenotypes in model species and humans (**Table 4**). Importantly, we provided evidence that dietary spermidine supplementation shifts these molecular ageing signatures towards a more favourable state in seawater-stage salmon, offering a promising strategy to mitigate associated health declines in aquaculture.

**Table 4.** Atlantic salmon metabolic ageing phenotypes mirror data in human and model mammals.

| **Metabolic ageing phenotypes observed in this study** | **Equivalent published phenotype in human or model rodents** |
| --- | --- |
| Lipid overload in SM (**Table 1**) | (Cree et al. 2004; Rivas, Morris, and Fielding 2011; Russ, Manickam, and Tipparaju 2025) |
| Phospholipid decline in SM (**Table 1**) | Pollard et al. 2017; Lee et al. 2020 |
| Metabolic flexibility in SM (**Figure 3**) | Conley, Jubrias, and Esselman 2000; Short et al. 2005; Kaczor et al. 2006; Lanza, Larsen, and Kent-Braun 2007 |
| Lysosomal dysfunction in VAT (**Figure 4, Supplementary Figure 6**) | No evidence in VAT, but strong evidence in ageing brain and adipose-derived stem cells from obese individuals (Burrinha et al. 2023; Wang et al. 2024) |
| Phagocytosis in VAT (**Figure 4, Supplementary Figure 6**) | Lu et al. 2021; Moss et al. 2023, 2024 |
| Inflammation and inflammaging in VAT (**Figure 4, Supplementary Figure 7**) | SM and VAT (Cornish and Cordingley 2024; Hayashi, Rakugi, and Morishita 2020; Kedlian et al. 2024; Kunz and Lanza 2023) |

Across different salmon life stages, lipidomics revealed a striking divergence between SM and VAT. Progression to the seawater ongoing stage (SWO) was associated with redistribution among TG species, a broad decline in multiple phospholipid species, and accumulation of total FAs in SM, whereas the FFA pool showed limited stage-associated changes. VAT in parr remained dominated by TGs and showed limited changes in total FAs and FFAs across life stages. These results indicate that VAT progressively loses its capacity to buffer lipid influx (from feed and/or circulating lipid), favouring lipid deposition in SM. This interpretation aligns with “adipose expandability/plasticity” models from mammals, where limitations in adipose storage capacity promote lipid spill over into non-adipose tissues, with downstream metabolic consequences (Frayn 2002; Virtue and Vidal-Puig 2010; Ou et al. 2022). In this context, the fibrosis/inflammation signature in the VAT proteome from SWO fish provides an additional mechanistic basis for the VAT–SM buffering model described above.

Functionally, the ageing phenotype we propose for muscle is consistent with reduced metabolic flexibility, defined as a diminished ability to shift substrate utilisation to meet energetic demand (Goodpaster and Sparks 2017; Jin et al. 2021; Kelley 2005). Compared to earlier life stages, we observed TG/FA accumulation together with lower abundance of free FFAs required as immediate substrates for mitochondrial β-oxidation and ATP generation, indicating that the expanded intramuscular TG/FA pool is not being efficiently mobilised. Taken with the observed proteome signature showing stage-associated increases in markers linked to FA uptake/trafficking and storage rather than oxidation, our data support a model where the muscle becomes increasingly TG/FA-loaded, yet less capable of mobilising this energy reservoir.

Beyond neutral lipid accumulation, SWO fish showed a reduction of phospholipid species (PC/PE classes), consistent with past work in SM (Jin et al. 2021). In mammals, mitochondrial membrane phospholipid composition is tightly linked to respiratory complex organisation, substrate oxidation capacity and overall mitochondrial efficiency; thus, phospholipid depletion or remodelling can contribute to reduced oxidative capacity and impaired metabolic switching (Decker and Funai 2024; Tasseva et al. 2013).

In VAT, the transition to SWO was marked by changes in lysosome and phagosome components, metabolic pathways and ECM–cytoskeleton proteins, indicating prominent remodelling of fat tissue during ageing in seawater. A key point emerging from the VAT proteome analysis was that smolts differ from SWO, not only in adipose structural remodelling, but also in immune lipid mediator pathways and innate defense mechanisms. Compared to smolt, SWO showed reduced abundance of proteins involved in arachidonic-acid/eicosanoid (oxylipin) and carbohydrate metabolism. This provides a plausible mechanism for interpreting post-smolt ageing as a state of immune/metabolic tuning, whereas SWO represents a state where tissue remodelling, lysosomal functions and innate defence capacity are comparatively reduced. Specifically, compared to parr and smolt, SWO-stage fish showed reduced abundance of multiple lysosomal hydrolases, components of acidification machinery, and proteins linked to lysosome–autophagosome fusion and autophagy regulation. Since lysosomal acidification is required for both degradative capacity (Lee et al. 2010; Mizushima and Komatsu 2011) and immune functions (including antigen processing) (McCoy 1990; Watts 2022; Yang et al. 2023), these coordinated changes suggest a general decline in endo-lysosomal competence in late-stage VAT.

In parallel, SWO VAT showed downregulation of proteins central to phagocytosis and innate immune defence. This is consistent with reduced pathogen recognition and killing capacity, and points to a potential weakening of adipose-resident innate immune functions in the late seawater phase (Trombetta et al. 2003). Importantly, VAT is increasingly recognised as an immunometabolic organ that switches between lipid storage/homeostasis and immune surveillance – a function reliant on intact lysosomal and phagocytic systems (Pignatelli et al. 2014; Simón et al. 2022). Thus, the decline observed in SWO VAT likely has dual consequences: reduced capacity for (i) lipid handling/recycling (including lysosome-dependent lipid recycling mechanisms) and (ii) immune defence and inflammatory regulation. It will be interesting to determine whether the reduced phagocytic capacity reflects age-related changes in the abundance and/or functional competence of phagocytic cells (e.g., macrophages, neutrophils and other granulocytes) (Schröder and Rink 2003), or indeed a reduced capacity to raise an effective immune response to pathogen challenges.

To test if the SWO metabolic ageing phenotype is modifiable, we supplemented seawater-stage salmon with dietary spermidine, a molecule known to enhance autophagy in model organisms and fish (Liao et al. 2021; Zhao et al. 2026). Spermidine supplementation not only enhanced autophagy, evidenced by changes in p62/SQSTM1 and ATG7 abundance in SM, it also remodelled SM lipid and protein signatures in a direction opposing the SWO ageing pattern. Specifically, vlcTGs decreased, phospholipids increased and FFAs increased, alongside increased abundance of markers indicating enhanced FA entry into mitochondria and oxidation (e.g., CPT1A and HADH) and increased glycolysis (e.g., PFKMA). These changes suggest spermidine promotes improved substrate handling and metabolic flexibility in SM, consistent with a liver study recently published in large yellow croaker (*Larimichthys crocea*) (Zhao et al. 2026). Interestingly, spermidine supplementation led to a series of proteomic changes indicating increased muscle health. This included increased abundance of multiple paralogous copies of XIRP2a, a protein that accumulates around regenerating muscle fibres in patients with SM myopathies (Nilsson et al. 2013) and POSTN, which has a role in muscle repair (Eulitz et al. 2013; Lees 2015). Conversely, spermidine downregulated LCP1, a protein associated with immune infiltration in humans (Pan et al. 2024) and T cell trafficking in mice (Joshi and Morley 2022) (**Figure 6g**).

In VAT, spermidine supplementation was associated with broader lipid-class changes, including increased abundance of several TG and phospholipid species, next to reductions in multiple lipotoxic lipid classes associated with inflammatory metabolic states. This pattern is consistent with a partial restoration of VAT’s role as a lipid “sink” and with reduced lipotoxic stress, aligning with our model that improving adipose storage capacity and lysosomal competence could reduce lipid spill over to muscle. While we were unable to obtain consistent VAT autophagy readouts due to the high lipid content of samples, the combined VAT lipidomic shifts and reduced inflammatory signature suggest spermidine acted to reprogram adipose lipid handling and inflammation.

Increased inflammatory responses have been demonstrated in the VAT of salmon fed on vegetable oil based diets (Xu et al. 2022). Our results show that this inflammatory signature is attenuated by spermidine, as reflected by reduced abundance of proteins linked to vascular activation and leukocyte trafficking, including ANGPT2A, CD99 and CD99L2 (Fiedler et al. 2006; Haugstøyl et al. 2023; Rutledge et al. 2022). CD99 and CD99L2 are heavily glycosylated transmembrane adhesion molecules that play key roles in leukocyte transendothelial migration, including monocyte movement across endothelial barriers. Notably, endothelial cells within VAT can acquire a pro-inflammatory, senescence-associated phenotype in obese humans (Pellegrinelli et al. 2014).

In addition, in VAT, spermidine enhanced protein abundance for known inhibitors of adipogenesis, including SLC43A3, a solute carrier transporter family member highly expressed in adipose tissues. Knockdown of SLC43A3 in adipocytes reduces both FA efflux, while increasing FA uptake and lipid droplet accumulation, while its overexpression decreases FA uptake in cells and decreases lipid droplet accumulation (Hasbargen et al. 2020). Spermidine also increased the abundance of AIFM2, a protein that translocates from lipid droplets to the outer mitochondrial inner membrane, where it supports glycolysis-driven thermogenesis by regenerating cytosolic NAD⁺ and supplying electrons to the electron transport chain under cold stress (Nguyen et al. 2020). Together, such changes are consistent with spermidine promoting a more lipolytic VAT environment in salmon—enhancing lipid mobilisation and utilisation, and thereby dampens inflammatory remodelling during the seawater grow-out phase.

## Conclusions

Overall, our data identify a coherent late-stage seawater phenotype in farmed salmon characterised by (i) impaired VAT lysosomal/phagocytic competence with fibrosis and inflammation, (ii) lipid deposition in muscle, and (iii) reduced muscle metabolic flexibility marked by lipid overload with limited FFA mobilisation and phospholipid depletion. Dietary spermidine supplementation showed that several of these phenotypes are modifiable, shifting lipid signatures and metabolic markers toward improved adipose buffering and muscle substrate utilisation. Together, these findings provide a case for farmed salmon offering a valuable comparative system for studying mechanisms of vertebrate ageing, while pointing towards nutritional strategies to improve health and welfare in commercial aquaculture.

## Limitations of the study

This study has limitations that caveat our interpretations and inform future work. Our life-stage comparisons were cross-sectional and confounded by commercial realities: parr were sampled from different facility/origins than smolt and SWO, while fish were fed standard commercial diets matched to life stage requirements, with dietary compositions therefore non-standardised. Further, the spermidine trial was relatively short (five weeks) and based on small sample sizes (n = 3 or 4). Though tank replication was included and adds confidence to our results, future studies should incorporate greater biological replication, longer interventions, and disease challenge endpoints to test if the inferred benefits of spermidine translate to enhanced resilience.

## Acknowledgements

We thank the Marine Environmental Research Laboratory, Machrihanish facility staff for their invaluable help with the sampling of fish from the spermidine feed trial.

## Data availability

All data referenced in the manuscript are included as supplementary files. The raw proteomics data will be submitted to the PRIDE repository. The raw lipidomics data will be submitted to Metabolights repository.

## Materials and Methods

### Ethics statement

Experimental protocols were performed following the Guidelines of the European Union (2010/63/EU) for the use of laboratory animals and following approval by the University of Stirling Animal Welfare and Ethical Review Board (AWERB 2023 15271 10615 and AWERB 2023 15246 11042). The feeding experiment was conducted in compliance with the Animals Scientific Procedures Act 1986 (Home Office Code of Practice, HMSO, London, January 1997) under project licence IFF90086F in accordance with EU regulation (EC Directive 86/609/EEC).

### Life-stage sampling

We sampled SM and VAT (n=6 each) from three Atlantic salmon life-stages (**Figure 1**): 1) parr aged 9 months (n=6, average weight of sampled fish: 99.6 ±18.1g); a juvenile freshwater stage, where the fish naturally inhabit streams/rivers and are held in recirculating systems or in cages or flowthrough tank systems in aquaculture; 2) smolt aged 9 months (n=6, average weight of sampled fish: 209.3 ± 42.9g), a more advanced stage of the salmon life-cycle, where the fish are physiologically adapting to life in seawater, primarily in response to photoperiod; and 3) seawater on-growing phase (‘SWO’) aged 20 months (n=6, average weight of sampled fish: 2,022 ± 0.4g); here, smolts that had completed smoltification were previously transferred into seawater pens for a growth phase leading to harvest at approximately 4-5kg weight. SWO represents approximately one year of ageing under commercial conditions from the sampled smolt stage.

Parr were sampled from the University of Stirling’s Niall Bromage Freshwater Research Unit (Origin: Benchmark, Iceland), while smolt and SWO were sampled from a commercial farm (MOWI, UK). All fish were fed standard commercial diets for each life stage (**Supplementary Table 1)**. Fish were euthanized by overdose by MS-222 (tricaine methanesulfonate; 300 mg/L; pH neutral) for 10 min, with death confirmed by destruction of the brain. Pure samples of VAT (around the intestine) and SM (from a standardised portion of the epaxial myotome taken just behind the dorsal fin) were dissected and snap-frozen on dry ice.

### Spermidine feeding experiment

The spermidine feeding experiment was conducted at the University of Stirling’s Marine Environmental Research Laboratory (MERL) (Machrihanish, Scotland). Atlantic salmon (outbred commercial aquaculture strain, eggs imported from Iceland, initial weight of 300 ± 0.02g) were kept in triplicate tanks (n=23 per tank). Salmon feed was formulated with and without the addition of 5mg/Kg spermidine (Sigma, S2626) and produced by Pontus Feed Technology Centre (Unit E, Hirwaun Industrial Estate, Hirwaun, Aberdare CF44 9UP). All diets contained the same concentration of fat, protein and carbohydrates. Details of feed formulation and nutritional profile are given in **Supplementary Figure 8**.

The experimental feeds were given in excess of expected appetite by autofeeders (ARVOTEC feeders) twice a day from 06:00-09:00 and 19:00-22:00 with uneaten feed collected to measure daily feed intake and calculate apparent feed intake. At the end of the trial all fish in each tank were killed at ∼6-12h post-feeding using the approach described above, then measured for weight and length (n=15 per tank). Next, VAT and SM from two fish per tank (n=6 per dietary treatment) were collected and snap-frozen on dry ice, using the same protocol as described above.

We also calculated several growth-associated parameters during the 5 week trial: 1) Feed conversion ratio (calculated as the dry weight of feed consumed divided by gain in wet weight; 2) Specific growth rate (calculated as: 100 x (lnWt – lnWo) x D -1, where Wo and Wt are the initial and end weights (tanks means, n=3) of the fish in a specific period, respectively, and D represents the number of feeding days; 3) Thermal growth coefficient (calculated as: 1000 x [(Wt(1/3) – Wo(1/3)) / °D], where Wo and Wt are the initial and end weights (tanks means, n=3) of the fish in a specific period, respectively, and °D represents degree-days, the sum of daily temperatures in °C in the specific period (or duration in days x average temperature in period); and 4) Weight gain (in g) (calculated as: Wt –Wo, where Wo and Wt are the initial and end weights (tanks means, n=3) of the fish in a specific period.

### Total fatty acid measurements

Total lipids were extracted from approximately 0.5 g of homogenised VAT or SM sample, using 20 volumes of ice-cold chloroform: methanol (2:1, v/v) according to (Folch, Lees, and Stanley 1957). Non-lipid contaminants were removed by washing with 0.88% (w/v) KCl, before total lipid content was determined gravimetrically after solvent evaporation under oxygen-free nitrogen followed by overnight desiccation under vacuum.

Fatty acid methyl esters (FAMEs) were prepared from total lipid extracts by acid-catalysed transmethylation at 50 °C for 16 h using 2 mL of 1% (v/v) sulphuric acid in methanol and 1 mL of toluene. FAMEs were then extracted and purified as described by (Tocher and Harvie 1988), based on the American Oil Chemists’ Society’s (AOCS) method for marine oils (AOCS 2007). FAMEs were separated and quantified by gas–liquid chromatography using a Fisons GC-8160 (Thermo Scientific, Milan, Italy) fitted with a 30 m × 0.32 mm i.d. × 0.25 μm ZB-wax column (Phenomenex, Cheshire, UK), using on-column injection and flame ionisation detection. Hydrogen was used as the carrier gas. The oven temperature was ramped from 50 °C to 150 °C at 40 °C min⁻¹ and then to 230 °C at 2 °C min⁻¹. Individual FAMEs were identified by comparison with known standards (Restek 20-FAME Marine Oil Standard, Thames Restek UK Ltd., Buckinghamshire, UK) and published reference data (Tocher and Harvie 1988). Data were acquired and processed using Chromcard for Windows (Version 1.19; Thermoquest Italia S.p.A., Milan, Italy).

Fatty acid (FA) content was quantified using heptadecanoic acid (17:0) as an internal standard added at a known concentration. Fatty acid concentration (FA; mg/ g lipid⁻¹) was calculated from the FA peak area relative to the internal standard, using the ratio of the internal standard concentration to its peak area. Lipid content was then used to express the results on a wet-weight basis (mg g sample⁻¹).

### Free fatty acid analysis

A specific sample amount was spiked with a mixture of deuterated internal standards, consisting of: C6:0-d11, C12:0-d23, C18:0-d35, C18:2-d4, C20:4-d11, C20:5-d5, C22:0-d43, C22:6-d5 and C24:0-d4 with 10 ng each (Cayman Chemical, Ann Arbor, USA). Methanol was added for protein precipitation and centrifuged. The clear solutions were analysed using an Agilent 1290 HPLC system with binary pump, multisampler and column thermostat with a Kinetex C-18, 2.1 x 150 mm, 2.7 µm column using a gradient solvent system of aqueous acetic acid (0.05 %) and acetonitrile. The flow rate was set at 0.4 mL/min, the injection volume was 1 µL. The HPLC was coupled with an Agilent 6470 Triplequad mass spectrometer (Agilent Technologies, Santa Clara, USA) with electrospray ionisation source. Analysis was performed with Multiple Reaction Monitoring in negative mode, with at least two mass transitions for each compound. All fatty acids were individual calibrated using Supelco 37-component FAME-Mix (Merck, Darmstadt, Germany) in relation to deuterated internal standards.

### Lipidomics data generation

Lipids were extracted from homogenised tissue samples following (Folch et al. 1957). Samples were extracted in a 2/1 (v/v) chloroform/methanol solution. The samples were left at 4 °C for 1 h then partitioned by addition of 0.1 M KCl. The mixture was centrifuged to facilitate phase separation. The lower chloroform layer was evaporated to dryness under a steady flow of nitrogen gas and reconstituted in methanol containing 5 mM ammonium formate.

Following extraction, lipids were analysed by liquid chromatography-mass spectrometry (LC-MS) using a Thermo QExactive Orbitrap mass spectrometer equipped with a heated electrospray ionization (HESI) probe and interfaced with a Dionex UltiMate 3000 RSLC system (Thermo Fisher Scientific, Hemel Hempstead, UK). Samples (10 μL) were injected onto a Thermo Hypersil Gold C18 column (2.1 mm × 100 mm; 1.9 μm) maintained at 50 °C. Mobile phase A consisted of water containing 10 mM ammonium formate and 0.1 % (v/v) formic acid. Mobile phase B consisted of a 90/10 (v/v) mixture of isopropanol:acetonitrile containing 10 mM ammonium formate and 0.1 % (v/v) formic acid. The LC gradient, maintained at a flow rate of 400 μL per min, was as follows; a starting condition of 65 % mobile phase A, 35 % mobile phase B. Mobile phase B increased from 35 to 65 % over 4 min, followed by 65 % to 100 % over 15 min, with a hold for 2 min before re-equilibration to the starting conditions over 6 min. All samples were analysed in positive and negative ionization modes over the mass-to-charge ratio (m/z) range of 250 to 2,000 at a resolution of 60,000.

The data was processed with Progenesis QI v2.4 software (Non-linear Dynamics). Relative fold quantification was performed using all ion normalization, with outputs limited to ion signals with intensities that showed 1.5-fold (or greater) difference between the study conditions and displayed statistical significance (p ≤ 0.05). This approach was performed for data acquired in both positive and negative ionization modes. Significant features were then identified using both Lipid Maps (Conroy et al. 2023) and the Human Metabolome Database (Wishart et al. 2022) with a mass error tolerance of 5 ppm. Confident annotations were allocated based on expected adducts for each lipid class versus the detected adducts.

Samples collected during the spermidine feeding trial were analysed by liquid chromatography-mass spectrometry (LC-MS) using a Waters Select Series Cyclic IMS mass spectrometer (cIMS) equipped with a heated electrospray ionization (Z-spray) probe and interfaced with a Waters Acquity Premier liquid chromatography system (Waters Corporation, Massachusetts, USA). All samples were analysed in positive and negative ionisation modes over the mass-to-charge ratio (m/z) range of 50 to 2,000 at a resolution of 60,000. The LC conditions and data analysis approach was as described above.

### Proteomics data generation

VAT and SM tissues were washed and resuspended in extraction buffer (5% sodium dodecyl sulfate (SDS), 50 mM triethylammonium bicarbonate (TEAB), pH 8.5) at a 1:10 (w/v) ratio and homogenized using a Precellys homogenizer (Bertin Technologies) at 5,000 g for 20 s in ceramic bead vials (Precellys Lysing Kit, CK Mix). Extracts were clarified by centrifugation (16,000 × g, 10 min), sonicated for 10 cycles (30 s on/off) on a Bioruptor Pico (Diagenode), and centrifuged again (16,000 × g, 10 min). Protein concentration was determined by bicinchoninic acid (BCA) assay.

Proteins (20 μg) were reduced and alkylated with 10 mM dithiothreitol (DTT) and 40 mM iodoacetamide (IAA) at 45 °C for 15 min, then processed on S-Trap micro columns (Protifi) following the manufacturer’s protocol with minor modifications. After acidification with phosphoric acid (2.5% final) and six-fold dilution in binding buffer (90% methanol, 100 mM TEAB), samples were loaded onto columns, washed three times with 150 μL binding buffer, and digested with 1 μg trypsin (Promega) in 50 mM TEAB at 47 °C for 2 h. Peptides were sequentially eluted with 50 mM TEAB, 0.2% formic acid, and 50% acetonitrile, pooled and dried by vacuum centrifugation.

Purified peptides were reconstituted in 0.1% formic acid and separated on an Aurora 25 cm × 75 μm C18 column (IonOpticks, Fitzroy, Australia) using a Thermo UltiMate 3000 RSLCnano system (Thermo Fisher Scientific) coupled online to a timsTOF fleX mass spectrometer (Bruker Daltonics, Bremen, Germany) equipped with a CaptiveSpray ion source. Peptide separation was achieved over a 70 min gradient at a flow rate of 200 nL/min with the column maintained at 50 °C. The trapped ion mobility spectrometry (TIMS) elution voltage was linearly calibrated to obtain reduced ion mobility (1/K0) values using three reference ions from the ESI-L Low Concentration Tuning Mix (Agilent Technologies, Santa Clara, CA, USA; m/z 622, 922, and 1222) in timsControl software (Bruker Daltonics).

Data were acquired in data-independent acquisition (DIA) mode using the Parallel Accumulation–Serial Fragmentation (diaPASEF) scheme (Meier et al. 2020). Full MS1 survey scans were recorded over the range of 100–1700 m/z with an ion mobility window spanning 1/K0 values from 0.65 to 1.45 Vs/cm². The diaPASEF method employed a defined set of precursor isolation windows tiled across the m/z and ion mobility dimensions to ensure comprehensive and unbiased sampling of all detectable peptide precursors within the surveyed range. Singly charged ions, which were fully resolved in the mobility dimension, were excluded from fragmentation. The collision energy was linearly ramped from 20 eV at 1/K0 = 0.6 to 59 eV at 1/K0 = 1.6.

Raw diaPASEF data files were processed directly in Spectronaut (version 19.2, Biognosys AG, Schlieren, Switzerland) using the directDIA workflow, which performs library-free analysis by generating a pseudo-spectral library from the DIA data and subsequently extracting quantitative information in a single integrated pipeline. Protein database searches were conducted against the Ensembl (release 115) annotation of the current Atlantic salmon reference genome (Ssal_v3.1; GCA_905237065.2), comprising 47,205 protein-coding genes. Trypsin/P was specified as the proteolytic enzyme with up to one missed cleavage permitted. Carbamidomethylation of cysteine (+57.0215 Da) was set as a fixed modification, while oxidation of methionine (+15.9949 Da) and deamidation of asparagine and glutamine (+0.9840 Da) were included as variable modifications. Precursor and fragment ion mass tolerances were determined dynamically by Spectronaut. Identification results were filtered at 1% false discovery rate (FDR) at both the precursor and protein levels using the target-decoy approach implemented in Spectronaut. Label-free quantification was performed using the MaxLFQ algorithm integrated within Spectronaut, with default settings for cross-run normalization and peptide-to-protein intensity rollup (Cox et al. 2014).

### Proteomics data analysis

Fold change and *p-values* for the obtained protein data were calculated using Sidoli’s method (Aguilan, Kulej, and Sidoli 2020). In short, where n was > 4 (for the life-stage samples), the Shapiro–Wilk test was first used to assess normality. For proteins where this test was significant, a non-parametric Mann Whitney test was applied. For proteins where non-significant (*p ≥* 0.05), the hypothesis of normality was accepted, and a parametric F test was applied to check for equal variances among the compared groups. If the F test was significant (*p* ≤ 0.05), a type 3 (two-sample unequal variance) T test was used, and if not significant *(p ≥* 0.05), a type 2 (two-sample) T test was applied. Fold changes were calculated as the ratio between the biological replicates between different life stages. *p* values were displayed as - log 2 p-values (> 4.3219 signifies *p* < 0.05). We did not apply correction for multiple testing as this approach has a high risk of generating false negatives in proteomics experiments (Pascovici et al. 2016). Where n < 4 (spermidine feeding experiment), the Shapiro-Wilks test was not applied, but all other calculations remained the same.

Volcano plots were generated using VolcaNoseR (Goedhart and Luijsterburg 2020). For pathway analysis, Ensembl identification number for proteins with fold change > 1.5 or < 0.6 were provided to ShinyGo 0.85 (Ge, Jung, and Yao 2020) as the foreground, while the entire set of proteins identified in the relevant experiment was provided as the background. g:Profiler (Raudvere et al. 2019) was used to acquire protein annotations by using the g:Convert tool for Atlantic salmon. Where there were no annotations available, we used orthologue annotations from zebrafish (*Danio rerio*) or mouse (*Mus musculus*) using g: Orth search.

### Immunoblotting assays

Muscle tissue was homogenised using a Rotor-Stator Homogenizer (Ultra-Turrax T10) in radioimmunoprecipitation assay (RIPA; Thermo Fisher Scientific) buffer containing protease inhibitors (Roche, Welwyn Garden City, UK). Protein concentrations were determined using the bicinchoninic acid (BCA) protein assay kit (Thermo Fischer Scientific.). A standard immunoblotting protocol was used with rabbit polyclonal antibodies against LC3 (1:3000, PM036), SQSTM1/p62 (1:500, Ab264313), and CPT1A (1:500, GeneTex, GTX636468). Band intensities were quantified using Fiji software. All blots were normalised to the estimated total protein concentration for each sample (Westerberg et al. 2025).

### Statistical analysis and data visualisation

For the life stage experiments, n=4 to 6 was used, while n=3 to 4 was used for the spermidine feeding experiments (biological replicates stated in figure legends). GraphPad Prism 10 was used for statistical analysis. To compare different FAs, free fatty acids (FFA) and lipidomics data, each was normalised by median. Unpaired two-tailed T tests were used to obtain *p* values for individual FA, FFA and lipidome (TGs etc). One-way ANOVA with Tukey’s test for post-hoc differences between pairs of stages was used to find significant differences in life stage data. *p <* 0.05 was considered a significant result. Heatmaps were generated using MetaboAnalyst *6.0* (Pang et al. 2024).

