## Supplementary Figures for "Spermidine enhances metabolic flexibility and attenuates inflammation associated with ageing in farmed Atlantic salmon"

Contains:

Supplementary Tables

Supplementary Figures

Supplementary Data legends

**Supplementary Table 1.** Details of commercial diet composition for parr, smolt and seawater ongoing salmon.

| Content | Units | Parr | Smolt | Seawater ongoing |
|---|---|---|---|---|
| Crude Protein | % of feed | 50 | 46.20 | 39.77 |
| Crude Fat | % of feed | 21 | 21.90 | 33.82 |
| Saturates | g/100g total lipids | No details found | 18.6 | 15 |
| Monoenes | g/100g total lipids | | 35.3 | 41.3 |
| n-3 PUFA | g/100g total lipids | | 17.9 | 14.5 |
| n-6 PUFA | g/100g total lipids | | 10.4 | 14.1 |
| EPA+DHA | g/100g total lipids | | 12 | 7.8 |
| Fish Oil | % of added oil | | 60.3 | 34.0 |
| Vegetable Oil | % of added oil | | 39.7 | 66 |

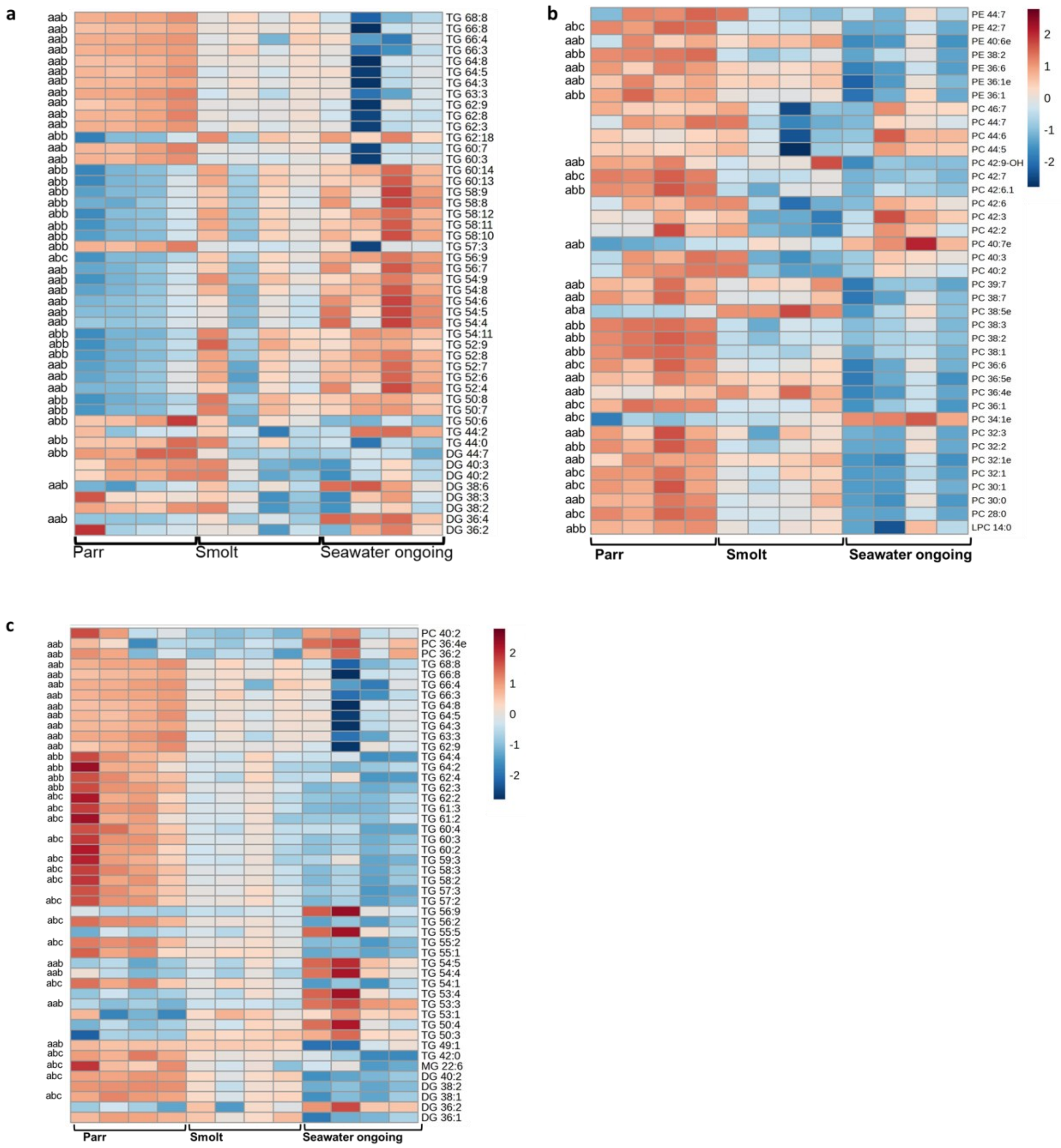


**Supplementary Figure 1.** Heatmaps to show abundance of lipids across three key life stages of salmon parr, smolt and seawater ongoing salmon. (a) Abundance of triglycerides and diglycerides in skeletal muscle (b) Abundance of phospholipids in skeletal muscle (c) Abundance of triglycerides, diglycerides and phospholipids in visceral adipose tissue.

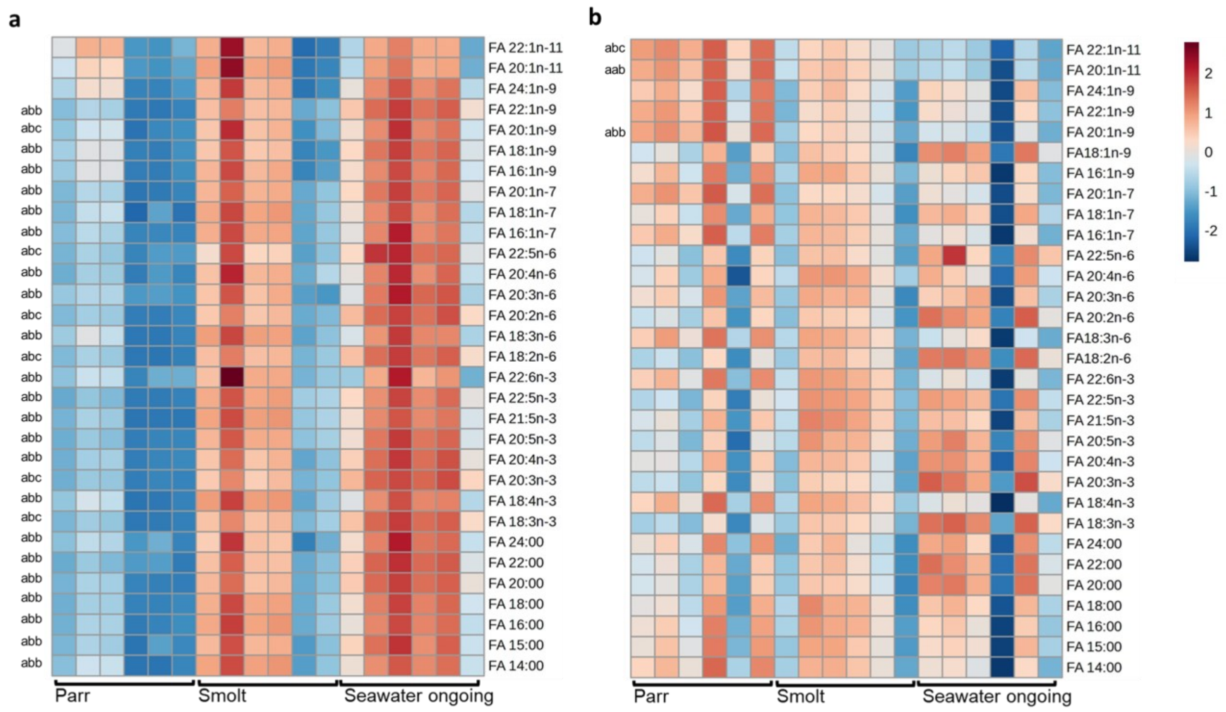


**Supplementary Figure 2.** Heatmaps to show abundance of fatty acids across three key life stages of salmon parr, smolt and seawater ongoing salmon. (a) skeletal muscle (b) visceral adipose tissue.

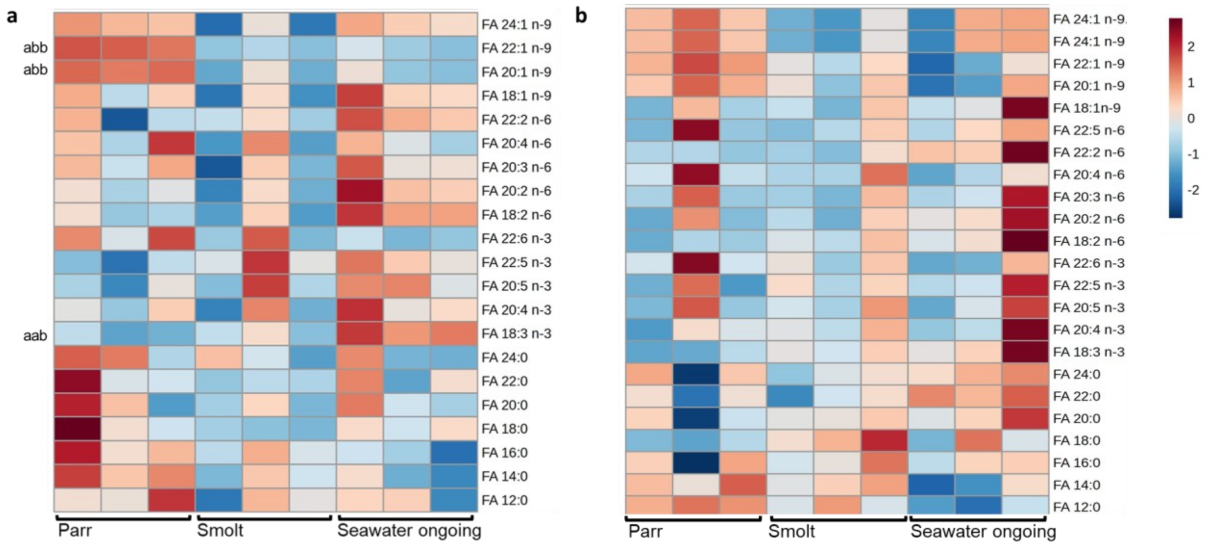


**Supplementary Figure 3.** Heatmaps to show abundance of free fatty acids across three key life stages of salmon parr, smolt and seawater ongoing salmon. (a) skeletal muscle (b) visceral adipose tissue.

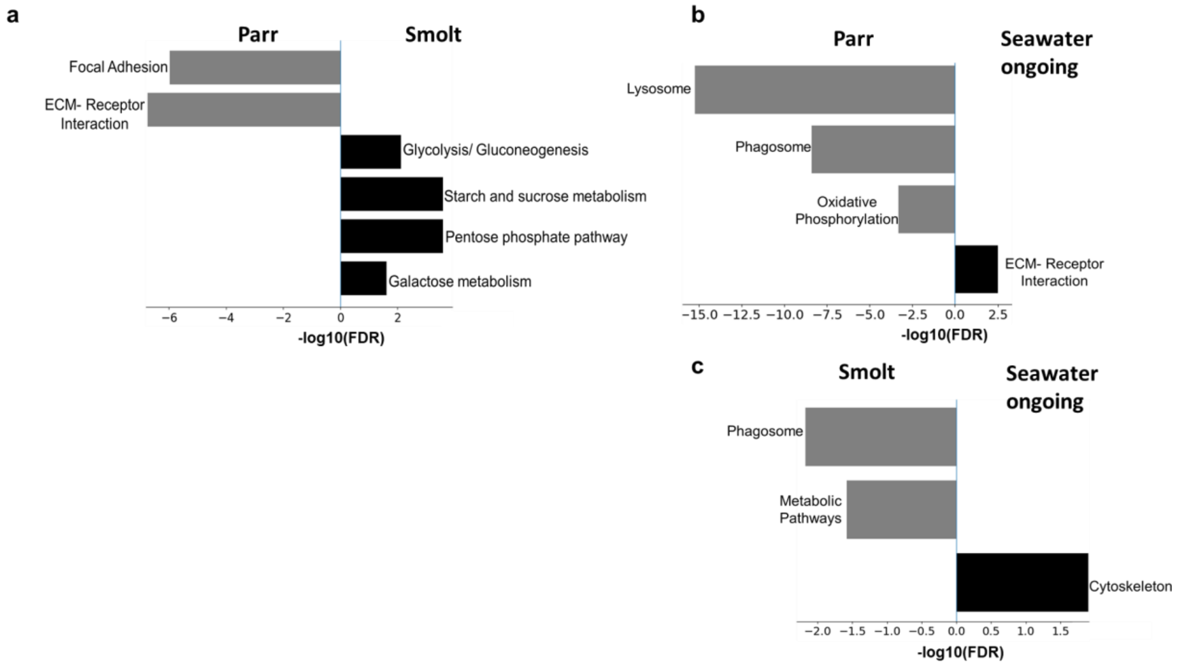


**Supplementary Figure 4.** GO term enrichment in the proteome data across three key life stages (parr, smolt and seawater ongoing salmon). (a) between parr and smolt in skeletal muscle (b) between parr and seawater ongoing in visceral adipose tissue (c) between smolt and seawater ongoing in visceral adipose tissue.

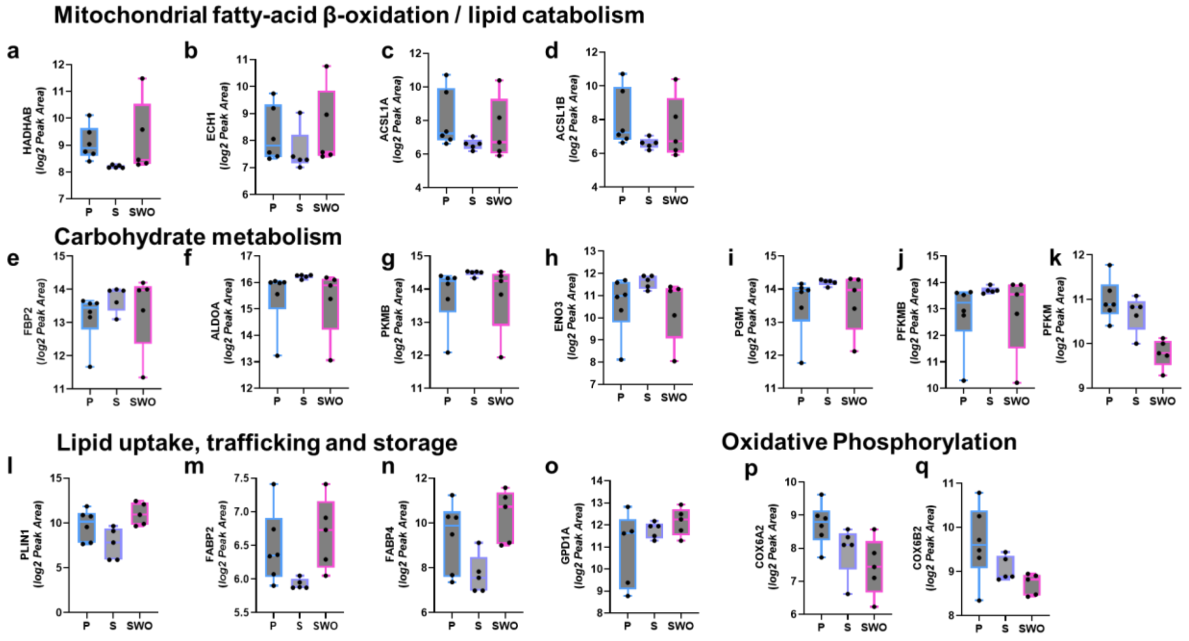


**Supplementary Figure 5.** Three-way box plot graphs comparing significant proteins from proteome analysis of skeletal muscle from parr, smolt and seawater ongoing salmon. (a)- (d) Mitochondrial β-oxidation/ lipid catabolism (e)- (k) carbohydrate metabolism (l)- (o) Lipid uptake, trafficking and storage (p)- (q) Oxidative phosphorylation.

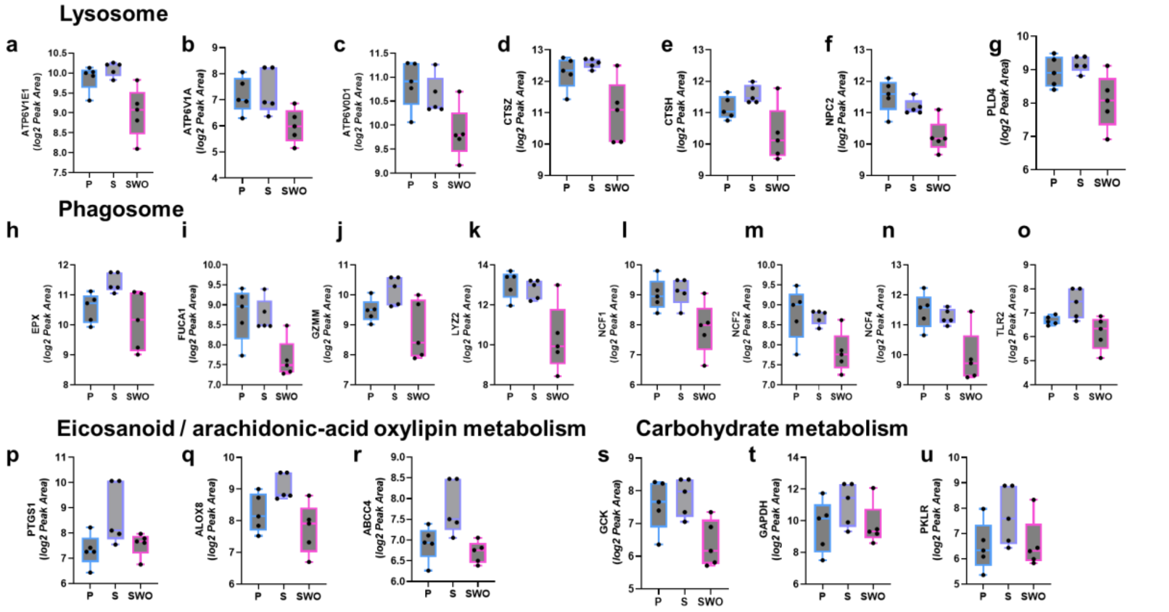


**Supplementary Figure 6**. Three-way box plot graphs comparing significant proteins from proteome analysis of visceral adipose tissue from parr, smolt and seawater ongoing salmon. (a)- (g) Lysosome (h)- (o) Phagosome (p)- (r) Eicosanoid/ arachidonic-acid oxylipin metabolism (s)- (u) carbohydrate metabolism.

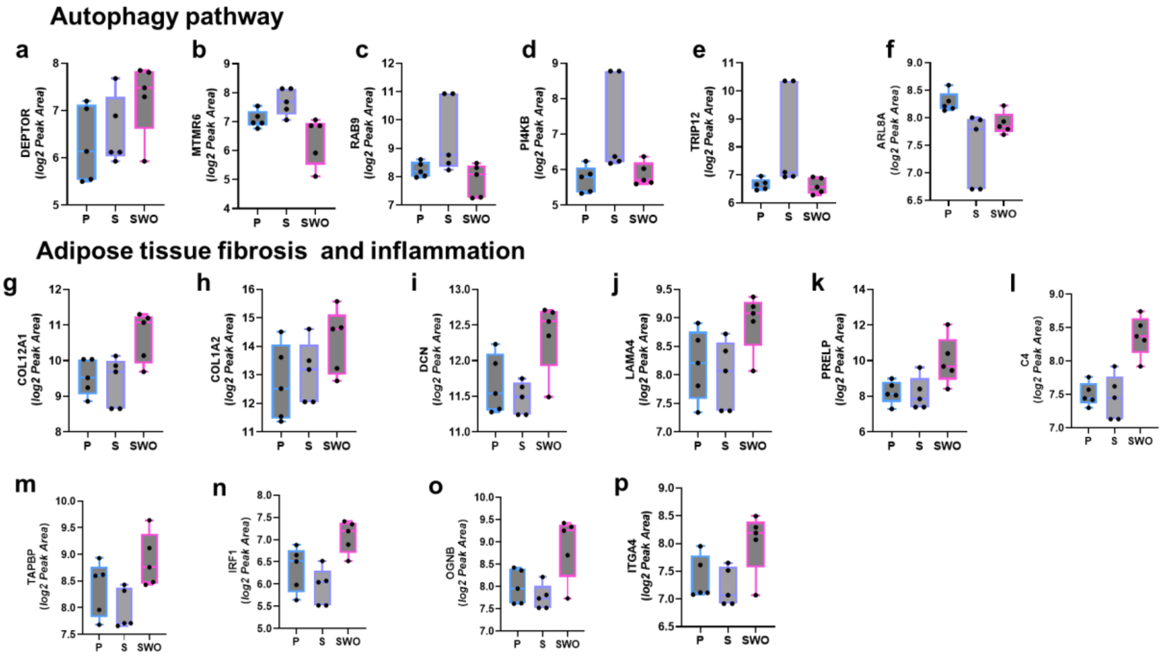


**Supplementary Figure 7**. Three-way box plot graphs comparing significant proteins from proteome analysis of visceral adipose tissue from parr, smolt and seawater ongoing salmon. (a)-(f) Autophagy pathway (g)- (p) Adipose tissue fibrosis and inflammation.

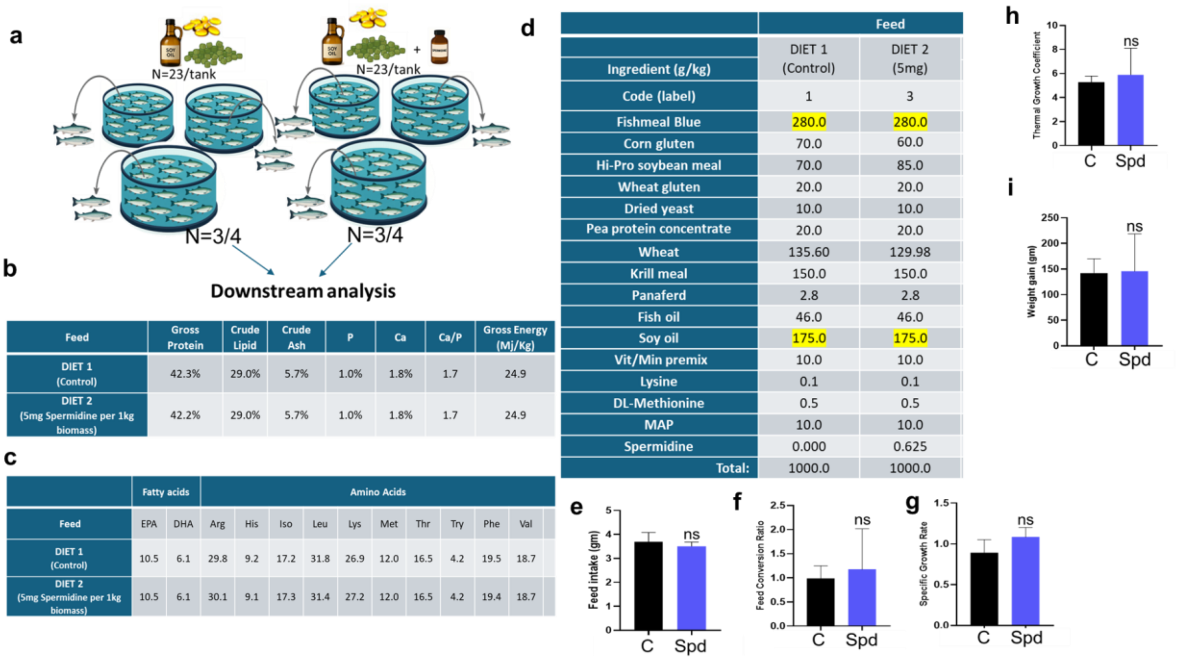


(b)

| Feed | Gross Protein | Crude Lipid | Crude Ash | P | Ca | Ca/P | Gross Energy (Mj/Kg) |
|---|---|---|---|---|---|---|---|
| DIET 1 (Control) | 42.3% | 29.0% | 5.7% | 1.0% | 1.8% | 1.7 | 24.9 |
| DIET 2 (5mg Spermidine per 1kg biomass) | 42.2% | 29.0% | 5.7% | 1.0% | 1.8% | 1.7 | 24.9 |

(c)

| | Fatty acids | | Amino Acids | | | | | | | | | |
|---|---|---|---|---|---|---|---|---|---|---|---|---|
| Feed | EPA | DHA | Arg | His | Iso | Leu | Lys | Met | Thr | Try | Phe | Val |
| DIET 1 (Control) | 10.5 | 6.1 | 29.8 | 9.2 | 17.2 | 31.8 | 26.9 | 12.0 | 16.5 | 4.2 | 19.5 | 18.7 |
| DIET 2 (5mg Spermidine per 1kg biomass) | 10.5 | 6.1 | 30.1 | 9.1 | 17.3 | 31.4 | 27.2 | 12.0 | 16.5 | 4.2 | 19.4 | 18.7 |

(d)

| | Feed | |
|---|---|---|
| Ingredient (g/kg) | DIET 1 (Control) | DIET 2 (5mg) |
| Code (label) | 1 | 3 |
| Fishmeal Blue | 280.0 | 280.0 |
| Corn gluten | 70.0 | 60.0 |
| Hi-Pro soybean meal | 70.0 | 85.0 |
| Wheat gluten | 20.0 | 20.0 |
| Dried yeast | 10.0 | 10.0 |
| Pea protein concentrate | 20.0 | 20.0 |
| Wheat | 135.60 | 129.98 |
| Krill meal | 150.0 | 150.0 |
| Panaferd | 2.8 | 2.8 |
| Fish oil | 46.0 | 46.0 |
| Soy oil | 175.0 | 175.0 |
| Vit/Min premix | 10.0 | 10.0 |
| Lysine | 0.1 | 0.1 |
| DL-Methionine | 0.5 | 0.5 |
| MAP | 10.0 | 10.0 |
| Spermidine | 0.000 | 0.625 |
| Total: | 1000.0 | 1000.0 |

**Supplementary Figure 8**. (a) Schematic diagram to show the details of spermidine feeding trial design. Showing complete nutrient profile of the formulated feed for Diet 1 (control) and Diet 2 (supplemented with spermidine 5 mg/kg biomass). (b) Gross protein, crude lipid and ash content of the diet along with Phosphorus, Calcium content (c) Total eicosapentaenoic acid (EPA), docosahexaenoic acid (DHA) and essential amino acid content. (d) Other ingredients including fishmeal, soyabean meal, fish oil, soy oil and spermidine. Quantification of basic growth parameters across 3 tanks with salmon fed on control diet and 5 mg/kg spermidine supplemented diet (N=3 each). (e) Feed intake (g) (f) Feed conversion ratio (g) Specific growth rate (h) Thermal growth coefficient (i) Weight gain (g).

**Supplementary Data 1a.** Complete skeletal muscle proteomics data set from parr (N=6), smolt (N=5) and seawater ongoing salmon (N=5) and its analysis.

**Supplementary Data 1b**. List of differentially expressed skeletal muscle proteome (*p<0.05*) between parr (N=6) and smolt (N=5) and smolt (N=5) and seawater ongoing salmon (N=5) and parr (N=6) and seawater ongoing salmon (N=5).

**Supplementary Data 2a**. Complete visceral adipose tissue proteomics data set from parr (N=5), smolt (N=5) and seawater ongoing salmon (N=5) and its analysis.

**Supplementary Data 2b.** List of differentially expressed visceral adipose tissue proteome (*p<0.05*) between parr (N=5) and smolt (N=5), smolt (N=5) and seawater ongoing salmon (N=5) and parr (N=5) and seawater ongoing salmon (N=5).

**Supplementary Data 3a**. Complete skeletal muscle proteomics data set from salmon fed on control (N=3) and 5mg/kg spermidine supplemented diet (N=3) and its analysis.

**Supplementary Data 3b**. List of differentially expressed skeletal muscle proteome ($p<0.05$) from salmon fed on control (N=3) and 5mg/kg spermidine supplemented diet (N=3).

**Supplementary Data 4a**. Complete visceral adipose tissue proteomics data set from salmon fed on control (N=3) and 5mg/kg spermidine supplemented diet (N=3) and its analysis.

**Supplementary Data 4b**. List of differentially expressed visceral adipose tissue proteome ($p<0.05$) from salmon fed on control (N=3) and 5mg/kg spermidine supplemented diet (N=3).
